# Cytokinin response regulator *ARR16* regulates seed coat permeability in Arabidopsis natural accessions

**DOI:** 10.1101/2023.02.15.528307

**Authors:** Naoto Sano, Frédéric Domergue, Helen M. North

## Abstract

The seed coat that encases the embryo is constituted from multiple specialized-cell layers and their permeability significantly influences seed quality traits that have major agronomic impact. The regulatory mechanisms that modulate seed coat permeability are, however, not well understood. Here, we identified a novel regulatory gene for seed permeability through a genome wide association study (GWAS) in Arabidopsis based on image analysis of tetrazolium staining. *Type-A ARABIDOPSIS RESPONSE REGULATOR 16* (*ARR16*) is a component of the signal transduction pathway for the plant hormone cytokinins (CKs), and in addition to less permeable seed coats, *arr16* mutant seeds were larger, had a longer lifespan, and more suberin phenolics, the hydrophobic lipid biopolyester components of cell walls that act as a water-repellant. Moreover, double mutants in CK receptor *ARABIDOPSIS HISTIDINE KINASE* genes, notably *ahk2 ahk4* and *ahk3 ahk4*, showed similar seed phenotypes to those of *arr16*. Based on naturally-occurring variation in the *ARR16* gene, eight haplotypes were detected and associated with permeable or impermeable phenotypes. Permeable haplotypes had significantly lower suberin autofluorescence compared to impermeable haplotypes. RT-qPCR analysis demonstrated that *ARR16* transcripts were highly abundant in developing seeds of representative accessions having permeable haplotypes but not in those of having impermeable haplotypes, indicating that these haplotypes were causal for *ARR16* transcript abundance and thereby regulate seed coat permeability in natural accessions. Our findings demonstrate a new role for CKs signaling in seed coat differentiation and that this underlies natural variation for seed permeability through the modulation of suberin accumulation.

**Significance Statement:** Seed coat permeability affects key traits that impact seed quality, such as dormancy, longevity, and germination tolerance to abiotic stress. Little is currently known about how seed coat permeability is modulated. Here, we developed an original method for quantification of seed coat permeability through imaging and used this in a genome-wide association study with Arabidopsis accessions. The CKs response regulator *ARR16* was identified as a causal gene thereby establishing a novel function for this phytohormone in seed coat differentiation. Moreover, this was linked to the modulation of suberin accumulation by *ARR16* and for the first time implicates CKs signal transduction in the control of suberin deposition in seeds.

## Introduction

The plant seed coat represents the interface through which the embryo interacts with the external environment and fulfills functions, such as protection, dispersal, controlling seed size and regulation of seed germination, dormancy and longevity^1^. Seed coats are formed of several overlying cell layers that accumulate large amounts of metabolites such as polyphenols, lignin, lignan, polysaccharides, cutin and suberin, whose properties impart physical and chemical resistance to the cells^2,3^. A number of mutants related to seed coat properties have been isolated and they often have altered seed coat permeability^4–7^. Natural variation in seed coat characteristics has also been widely documented and can be exploited in genome wide association studies (GWAS). Here, we identified type-A ARABIDOPSIS RESPONSE REGULATOR 16 (ARR16) as a novel regulator of seed permeability through GWAS based on image analysis of tetrazolium staining. *ARR16* is a component of the signal transduction pathway for the plant hormone cytokinins (CKs)^8^, and functional analysis of mutants revealed that *ARR16* modulates the accumulation of suberin phenolics.

## Results and Discussion

A group of Arabidopsis mutants, collectively called “*transparent testa*” (*tt*), have altered seed coat properties as they are defective for the production of seed coat pigments, proanthocyanidins (PAs) that are accumulated in the innermost integumentary layer of the seed coat^3^. More than 20 *tt* mutants have been reported, and their seeds can be characterized by tetrazolium (TZ) staining as PAs decrease seed coat permeability^4^. The TZ salts are metabolically reduced to red colored formazans by endogenous enzymes, such as NADH-dependent reductases, when they are assimilated by live cells. After TZ staining, seeds of *tt* mutants, such as *tt4*, appear redder compared to wild type (WT) due to the increased permeability of their seed coats (Fig. 1a). *tt* mutants exhibit additional seed phenotypes such as lower dormancy, shorter longevity and reduced tolerance of germination to toxic drugs^4,10^, emphasizing the importance of permeability for seed quality. Although, seed coat defects in the maternally deposited endosperm cuticle and tannic cell walls of *tt* seeds were recently reported^11,12^, our current knowledge regarding the regulation of seed coat permeability remains fragmentary. Notably, to date no genetic screen has been performed in Arabidopsis that specifically assesses seed coat permeability by TZ staining.

**Figure 1.**
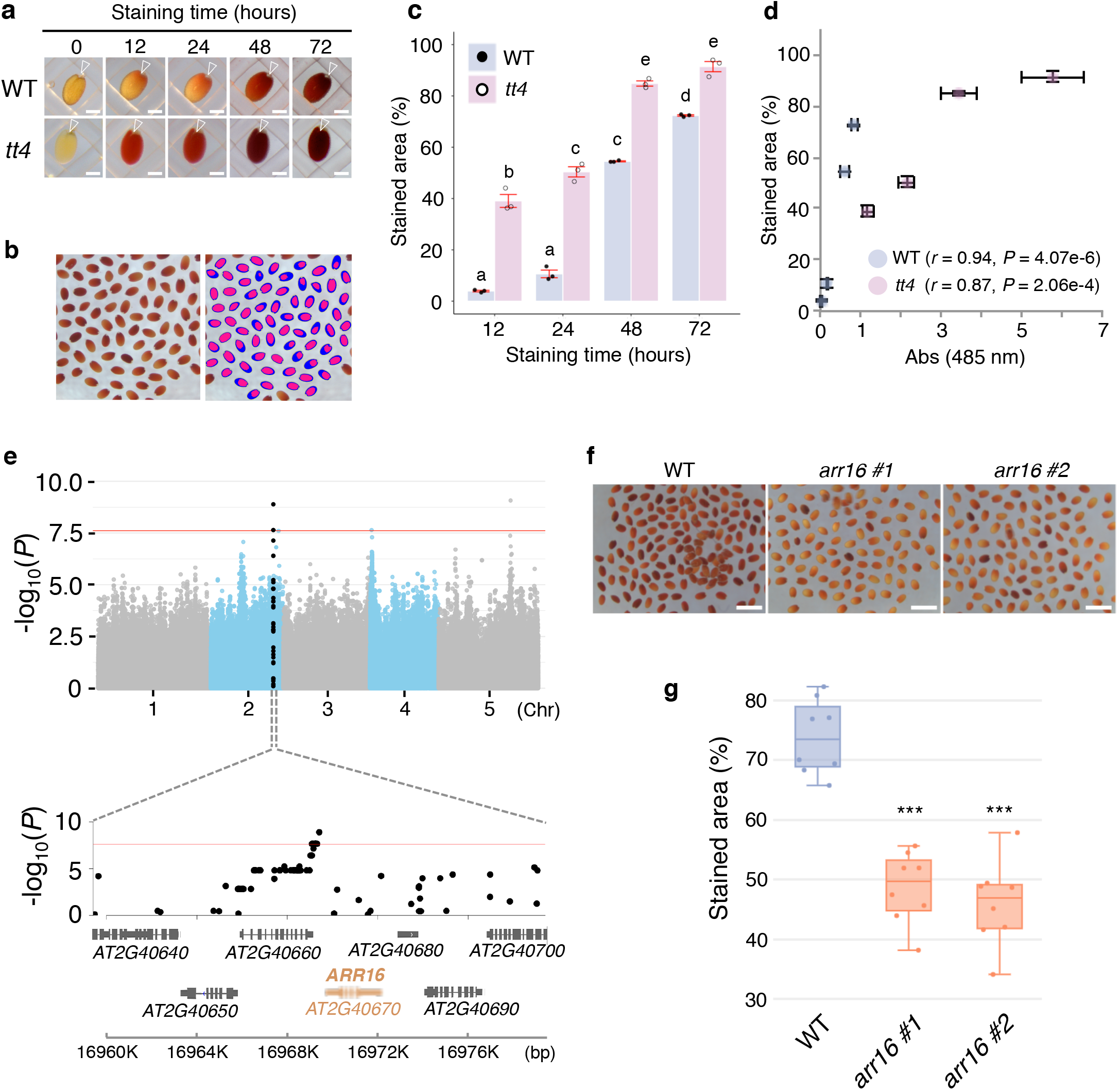
Identification of *ARR16* as a negative regulator of seed coat permeability. **a**, Representative seeds of WT and *tt4* at 0, 12, 24, 48 and 72 hours after tetrazolium (TZ) staining. White arrowhead indicates the position of the hilum. Scale bars represent 200 μm. **b**, Quantification of seed coat permeability by image analysis (left; original image, right; overlay of binarized images). Blue and pink regions indicate total seed area and total stained area with TZ solution, respectively. The detailed method is described in Supplementary Fig. 1. **c**, Bar graphs with jitter points showing TZ stained area (%) for WT and *tt4* seeds quantified by image analysis. Values presented are means, red error bars are ±SE (n=3). Different letters indicate significant differences (*P* < 0.05, Tukey–Kramer tests). **d**, Scatter plot of the relationship between the stained area (%) and formazan production measured by absorbance (Abs) at 485 nm in staining seeds of WT and *tt4* with TZ solution. Values presented are means, error bars are ±SE (n=3) (*r* = Pearson’s correlation coefficient, *P* = p-values for the correlation). **e**, Genome-wide association study of seed coat permeability to TZ. Manhattan plot of GWAS results for stained area (%) of seeds at 72 hours after TZ staining (top). SNPs within ±10 kb of the major SNP peak on chromosome 2 are shown as black dots. Enlarged view of SNPs and loci within ±10 kb of the major SNP peak on chromosome 2 (bottom). The red horizontal lines indicate a statistically significant threshold for the −log10(*P*) (FDR < 0.01, Benjamini– Hochberg method). **f**, Seeds of WT and *arr16* mutants at 72 hours after TZ staining. Scale bars represent 1 mm. **g**, Boxplot with jitter points (n=8) showing stained area (%) of WT and *arr16* mutant seeds at 72 hours after TZ staining (\*\*\**P* < 0.001, Dunnett’s test).

The hilum is the attachment point of the seed to the funiculus, the structure connecting the maternal plant to the seed during seed development. Many studies have designated the hilum as the main entry path for water during imbibition of matured dry seeds^13,14^. Similarly, during TZ staining, red colored formazan spreads from the hilum to the whole of the seed (Fig. 1a). An image analysis approach enables the accurate analysis of many samples with fewer operating procedures in a short time. Thus, we developed an image analysis system that can quantify seed coat permeability (%) following establishment of a color threshold and binarizing image of the TZ stained seeds (Fig. 1b) (Supplementary Fig. 1). This image analysis confirmed that the permeability (stained area%) of *tt4* is significantly higher than that of WT during TZ staining (Fig. 1c) and values obtained by the image analysis were significantly and positively correlated with those obtained measuring absorbance (Fig. 1d), which is a traditional quantification method for TZ staining. We next performed a GWAS for seed coat permeability based on our image analysis phenotypes with seed lots from 145 Arabidopsis accessions (Supplementary Table 1). At 48 and 72 hours after TZ staining, a wide phenotypic variation in staining percentage was observed (Supplementary Fig. 2). Fifteen single nucleotide polymorphisms (SNPs) were detected that significantly correlated with the stained area (%) at 72 hours after TZ staining (Supplementary Table 2). Among them, 10 SNPs were located in the promoter region of *ARR16 (AT2G40670*) on chromosome #2 (Fig. 1e). A similar peak of SNPs was observed in the dataset at 48 hours after TZ staining (data not shown). Seeds from two loss-of-function mutation lines *arr16 #1* and *arr16 #2* were significantly less permeable than those of WT when subjected to TZ staining (Fig. 1f,g). Similar phenotypes were observed in two T-DNA insertion lines having knockdown effects on *ARR16* expression during seed development (Supplementary Fig. 3), indicating that *ARR16* is a causal gene in our GWAS and negatively regulates seed coat permeability. Expression profile data accessible in a public microarray database indicate that transcript levels for *ARR16* are highest during late seed coat development and are also present in the developing endosperm (Supplementary Fig. 4a). We confirmed that the transcript is most abundant at the mature green phase of seed development (about 15 days after flowering (DAP)) in developing seeds by RT-qPCR analysis (Supplementary Fig. 4b). Moreover, F1 seeds from reciprocal crosses between WT and *arr16* confirmed that the genotype of the maternal seed coat tissue defined the permeability phenotype (Supplementary Fig. 5). These results demonstrate that *ARR16* expression modulates seed coat permeability, probably during the development of the seed coat. Additionally, seed size and weight were higher in *arr16* compared to WT (Fig. 2a,b). Moreover, *arr16* seeds showed better vigor (T_50CG_) and longevity (P_50_) than WT after controlled deterioration treatment (CDT), an artificial accelerated aging treatment (Fig. 2c,d; Supplementary Fig. 6a,b).

**Figure 2.**
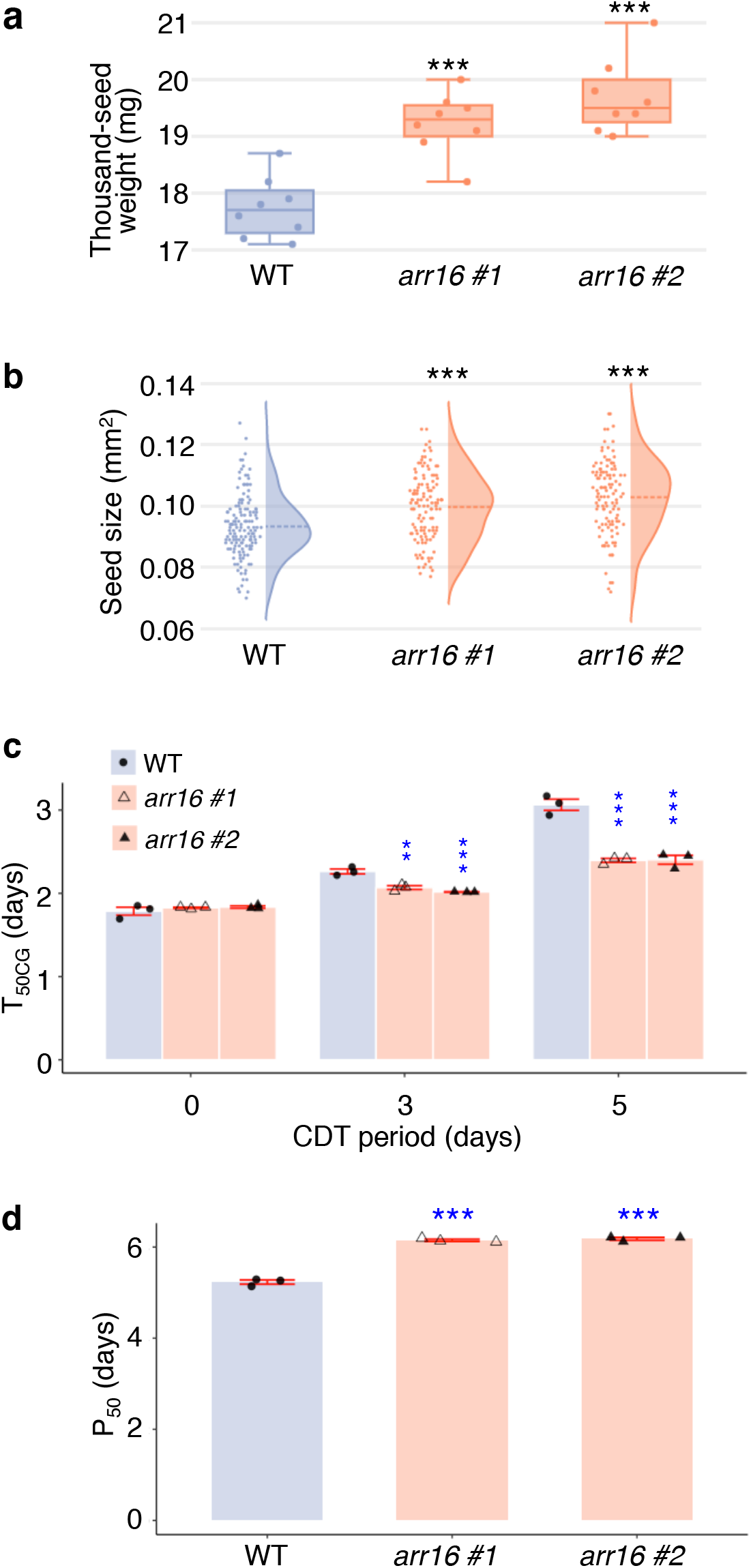
Contribution of *ARR16* to seed traits. **a**, Boxplot with jitter points (n=8) showing thousand-seed weight of WT and *arr16* mutant seeds (\*\*\**P* < 0.001, Dunnett’s test). **b**, Split violin plots with rotated kernel density estimates showing seed size of WT and *arr16* mutants (\*\*\**P* < 0.001, Dunnett’s test). Jitter points represent seed size data (n=143, 126, and 117 for WT, *arr16 #1* and *arr16 #2*, respectively). Dashed line shows the mean value for each genotype. **c**, Seed vigor of WT and *arr16* mutants after controlled deterioration treatment (CDT). Bar graphs with jitter points showing cotyledon greening speed T_50CG_ (time required for 50% of viable seeds to develop green cotyledons) was analyzed at 0, 3 and 5 days after CDT. Values are means, red error bars are SE (n=3), and blue asterisks indicate significant differences (\*\**P* < 0.01, \*\*\**P* < 0.001, Dunnett’s test). **d**, Seed longevity of WT and *arr16* mutants. Bar graphs with jitter points indicating index of seed longevity P_50_ (time for survival rate to decrease to 50%) during CDT. Values presented are means, red error bars are SE (n=3) and blue asterisks indicate significant differences (\*\*\**P* < 0.001, Dunnett’s test).

Among the components of the seed coat known to affect permeability, suberin, a lipid and phenolic cell wall heteropolymer, is mainly accumulated in the hilum region of seed coat and can be detected by UV autofluorescence (AF)^5, 15, 16^. More suberin AF was observed in the hilum of *arr16* than WT (Fig. 3a) and a significant difference in the level of AF signal was confirmed by image analysis (Fig. 3b). Stronger staining of the hilum area of the seed coat in *arr16* compared to WT, was also observed using the lipophilic dye Sudan Red 7B suggesting that the accumulation of suberin is higher in *arr16* (Supplementary Fig. 7). Additionally, GWAS based on hilum AF measured by image analysis was performed and resulting SNPs associated with hilum AF level were compared with those associated with TZ stained area (Fig. 3c). Interestingly, the 10 SNPs located in the promoter of *ARR16* (Supplementary Table 2) were significantly associated with both traits. This suggests that *ARR16* regulates the variation of permeability in natural accessions, at least partially, by affecting suberin deposition in the hilum. Analysis of seeds for lipid polyester monomer composition revealed that certain dicarboxylic acids (DCAs) and phenolics were accumulated to significantly higher levels in *arr16* compared to WT (Fig. 3d), while changes in other suberin aliphatics were minor (Supplementary Table 3). The strongest alterations in DCAs were observed for 18:1-DCA and 24:0-DCA, which are specific suberin markers (Fig. 3e). A major difference was observed for methyl 3,4 dihydroxy benzoate levels, which were nearly two-fold higher in *arr16* than WT (Fig. 3f). This small phenolic could be a constituent or intermediate of the phenolic domain of suberin. This corroborates the observed differences in suberin AF as phenolic rings and conjugated bonds in suberin will contribute to overall fluorescence^17^. RT-qPCR analysis for suberin biosynthesis genes was performed using RNA from developing seeds and differential gene expression patterns were observed between *arr16* and WT, such as for *FACT* (Supplementary Fig. 8). These suggest that the normal expression pattern of genes related to suberin biosynthesis may be disrupted in *arr16*, resulting in higher accumulation of DCAs and suberin phenolics in the mutant.

**Figure 3.**
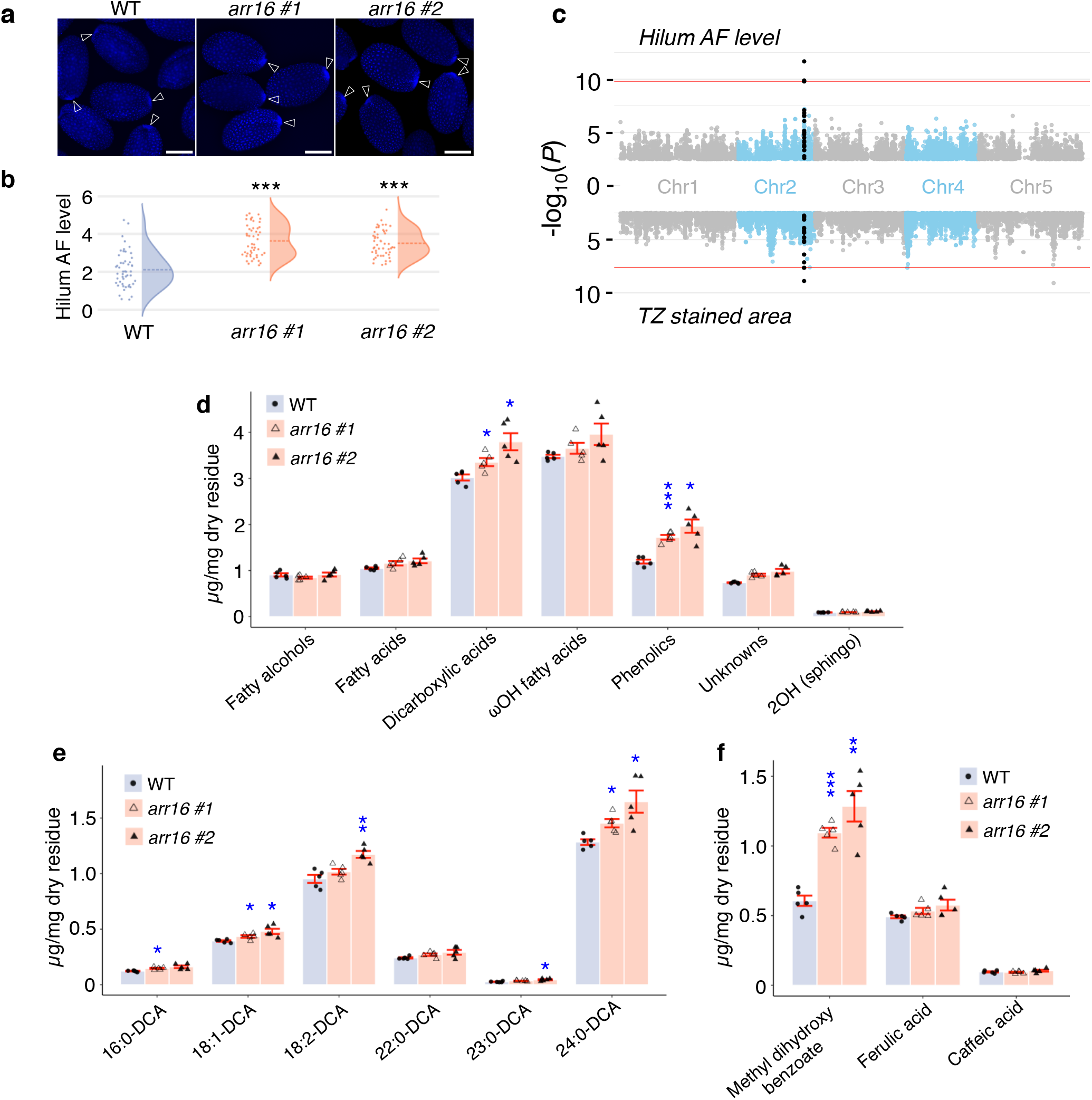
*ARR16* regulates suberin accumulation in seeds. **a**, Dry seeds of WT and *arr16* mutants under UV light. White arrowhead indicates the position of the hilum. Scale bars represent 200 μm. **b**, Split violin plots with rotated kernel density estimates (dashed line indicates mean value) and jitter points (n=50) showing the autofluorescence (AF) level in hilum region of seeds in WT and *arr16* mutants (\*\*\**P* < 0.001, Dunnett’s test). **c**, Genome-wide association study for suberin fluorescence in the hilum and seed coat permeability. Miami plot of GWAS results for hilum AF level of dry seeds (top) and stained area (%) of seeds at 72 hours after TZ staining (bottom). SNPs within ±10 kb of the major SNP peak on chromosome 2 are shown as black dots. The red horizontal lines indicate a statistically significant threshold for the −log_10_(*P*) (FDR < 0.01, Benjamini–Hochberg method). **d**, Substance class composition of suberin aliphatics, phenolics, unknowns and sphingolipids in WT and *arr16* mutant seeds. Bar graphs with jitter points indicating total amount of each compound class per a dry weight basis of delipidated material residue. Values presented are means and red error bars are SE (n=5) and blue asterisks indicate significant differences (\**P* < 0.05, \*\*\**P* < 0.001, Dunnett’s test). **e**, Detailed composition of dicarboxylic acids (DCA) and **f,** phenolics in WT and *arr16* mutant seeds. Bar graphs with jitter points indicating amount of each monomer with respect to dry weight of delipidated material residue. Values presented are means, red error bars are ±SE (n=5) and blue asterisks indicate significant differences (\**P* < 0.05, \*\**P* < 0.01, \*\*\**P* < 0.001, Dunnett’s test).

The naturally-occurring variation in *ARR16* gene sequence were characterized further through a haploblock analysis. Based on polymorphism information available for 1135 accessions, a total of 17 SNPs were detected in *ARR16* and three major haploblocks with strong linkage disequilibrium (LD) were identified (Fig. 4a). The three haploblocks consist of eight haplotypes, with the reference accession Col-0 classified as haplotype 1 (H1) (Fig. 4b). We then reanalyzed data of TZ stained area for 145 accessions based on these haplotypes, except for H6, which has the lowest frequency in Arabidopsis accessions and is not included in our data set. H1, H2 and H3 could be classed as permeable type haplotypes and H4, H5, H7 and H8 as impermeable ones, because no significant difference was observed for accessions having the former haplotypes, compared to reference H1 accessions, while accessions with the latter haplotypes showed remarkably lower permeability (Fig. 4c). Similarly, the permeable type haplotypes had significantly lower hilum AF level compared to impermeable ones (Fig. 4d). RT-qPCR analysis demonstrated that *ARR16* transcripts were highly abundant in developing seeds of representative accessions in the permeable type haplotypes but not in those of the impermeable type haplotypes, except for one accession in H8, Di-G, indicating that these haplotypes affect *ARR16* transcript levels and thereby regulate seed coat permeability in natural accessions (Fig. 4e). Genetic variation in the UTR could affect the expression level and stability of transcripts, for example by modifying regulatory elements involved in the interaction of the UTR with proteins and miRNAs^18^. To investigate this possibility, we carried out *in silico* analysis of the effect of *ARR16* haplotypes on mRNA secondary structure, but no significant structural changes were predicated (Supplementary Figs. 9 and 10). The causal SNPs affecting *ARR16* mRNA steady-state levels in natural accessions are most likely, therefore, to be located in haploblocks 1 and 2, which are in the promoter. While the association of *ARR16* haplotypes with natural variation for seed permeability may suggest local selection as this affects seed traits that impact plant fitness, no geographical or environmental links between accessions in a given class were evident.

**Figure 4.**
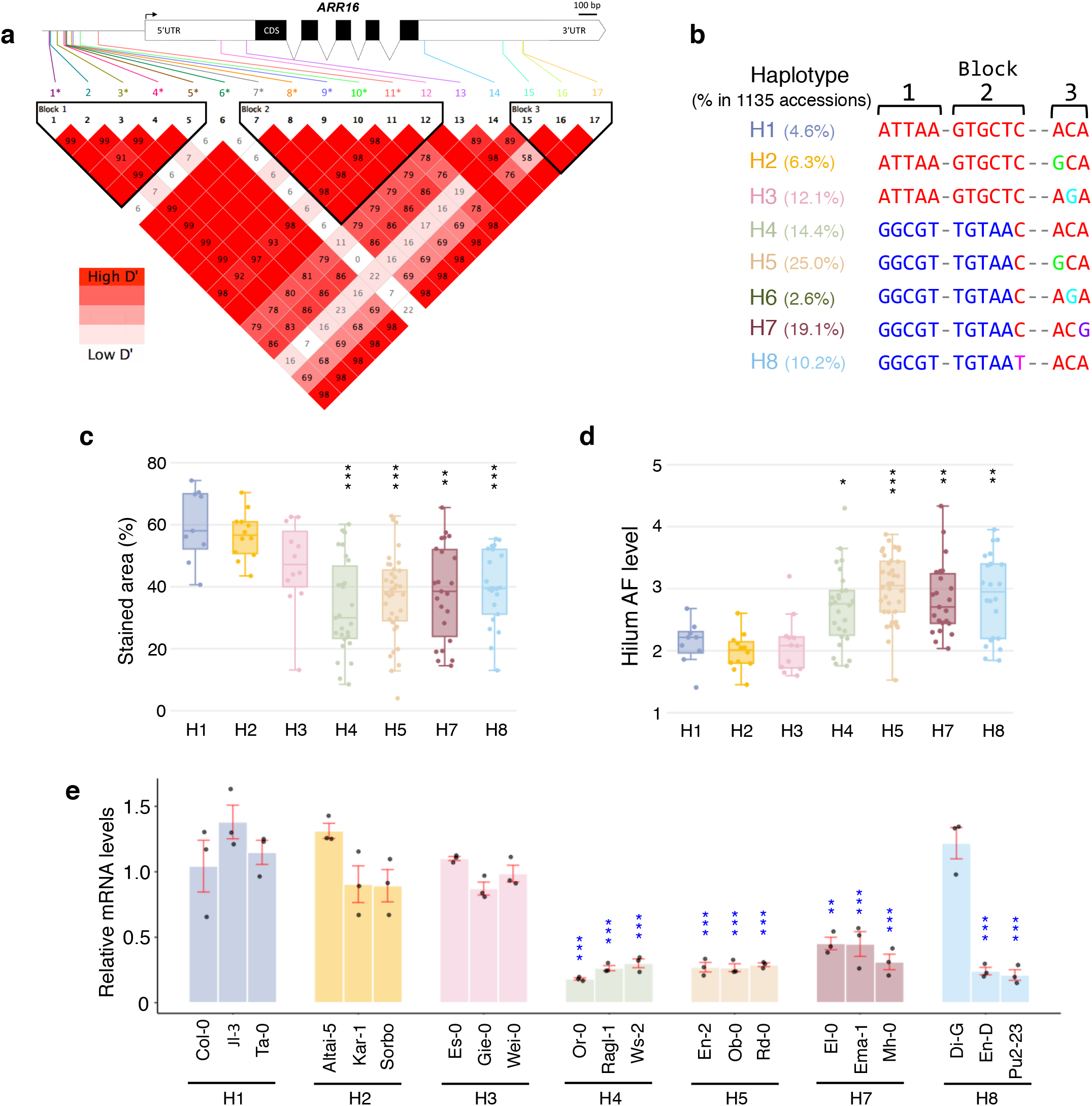
*ARR16* haplotypes and their involvement in seed coat permeability. **a**, Natural variants of *ARR16* in Arabidopsis. Schematic representation of the positions of 17 SNPs in the *ARR16* gene (top). UTR; untranslated region, CDS; coding sequence, diagonal lines; introns. Scale bar represents 100 base pairs. SNP numbers with an asterisk are significantly correlated to seed coat permeability (TZ staining area) in the GWAS. Linkage disequilibrium (LD) analysis of the 17 SNPs in *ARR16* gene (bottom). The D’ value, index of LD of each SNP pair, was obtained using Haploview software and is shown in the squares as D’× 100 (no value is indicated for squares with a D’ value of 1). Haploblocks were constructed with the default algorithm of Haploview (95% confidence intervals of the D’ values). **b**, Based on the three haploblocks, eight haplotypes (H1, H2, H3, H4, H5, H6, H7 and H8) can be defined for *ARR16*. The letters under the three haploblocks indicate the 5, 6 and 3 SNP alleles contained in each, respectively. The numbers in brackets indicate the frequency of each haplotype among the 1,135 Arabidopsis accessions. Boxplots of stained area (%) of seeds at 72 hours after tetrazolium staining (**c**) and the autofluorescence (AF) level in the hilum region of dry seeds (**d**) for each haplotype of *ARR16 (*P* < 0.05, \*\**P* < 0.01, \*\*\**P* < 0.001, Dunnett’s test, compared to H1). Jitter points represent each value for the stained area (%) (n=9, 12, 12, 27, 36, 23 and 22 for H1, H2, H3, H4, H5, H7 and H8, respectively). **e**, Expression levels of *ARR16* in developing seeds for each haplotype. Bar graphs with jitter points indicating *ARR16* expression in developing seeds at 15 DAP analyzed by RT-qPCR. Values are means, red error bars ±SE (n=3), and significant differences are indicated with blue asterisks (\*\**P* < 0.01, \*\*\**P* < 0.001, Dunnett’s test, compared to Col-0).

Three CK receptors (histidine kinases; AHK2, 3, 4) have been identified in Arabidopsis and the triple mutant *ahk2/3/4* was reported to have larger seeds^19^. Although this phenotype was suggested to be under the control of maternal or endosperm tissue, the potential role for CKs in the seed coat was not examined further. We confirmed that *ahk* double mutants also produce larger and heavier seeds than WT (Supplementary Fig. 11). The effect of CKs on seed size has been reviewed in detail^20,21^, and CKs seem to be positively correlated with seed size based on the phenotypes of mutants in CKs biosynthesis and metabolism, thus contrasting with *ahk* phenotypes. It should be noted that the *ahk2/3/4* mutant was dwarf and developed few seeds, and that seed mass is generally negatively correlated with total seed yield. Nevertheless, *arr16* plant growth was equivalent to WT (data not shown), suggesting that *ARR16* may regulate seed size through a seed-specific CKs signaling pathway. Arabidopsis has 11 type-B ARRs and 12 type-A ARRs, which are functionally redundant and have similar expression patterns^22,23^. As the single *arr16* mutant had clear seed phenotypes and our GWAS did not detect significant SNPs in loci related to other CKs signaling genes, *ARR16* may have a non-redundant role in CKs signaling in developing seeds. Moreover, *ahk* double mutants, especially *ahk2/4* and *ahk3/4*, showed similar seed phenotypes to those of *arr16* in terms of permeability, longevity and suberin accumulation (Supplementary Figs 12, 13 and 14). These suggest that CKs signaling itself may have a negative effect on suberin deposition, thereby regulating seed coat permeability and longevity. Type-A ARRs, to which ARR16 belongs, act as negative-feedback regulators of CKs signaling^8^, although some of them regulate downstream CKs responses directly. For example, type-A ARR4, 5 and 6 were shown to interact with ABSCISIC ACID (ABA) INSENSITIVE 5 and thereby negatively control ABA signaling in seed germination^24^. Moreover, ABA has been shown to positively regulate suberin formation in roots^25^, in agreement with our results linking *ARR16* expression to suberin production. Identifying other factors that interact with ARR16 in seed coat cells to transmit CKs signals downstream may clarify this enigma of *arr16* and *ahk* exhibiting similar modifications in seed coat phenotypes. Several transcription factors are already known that modulate suberin deposition in seeds^26–28^, however, the implication of a CKs signal transduction factor has not previously been reported. Intriguingly, during differentiation of the xylem pole in roots, CKs signaling is reduced to produce unsuberized ‘passage cells’^29^, indicating that the level of CKs signaling determines the degree of plant cell suberization. Our findings provide important evidence of an additional role for CKs signaling in seed coat differentiation in addition to controlling the suberization of specific cells in plant tissues.

## Materials and Methods

### Plant materials

Homozygous *tt4* mutant (SALK_020583) in Col-0 accession background was confirmed by PCR using primers designed according to the SIGnAL (Salk Institute Genomic Analysis Laboratory) website (http://signal.salk.edu/tdnaprimers.2.html). Col-0 (CS39288) and 144 natural accessions (CS76636) in Arabidopsis were obtained from the Nottingham Arabidopsis Stock Centre (NASC) (http://arabidopsis.info/). The *arr16* mutant lines (*arr16 #1* and *arr16 #2*) were generated using CRISPR/Cas9-mediated genome editing in Col-0 background as described previously^30^. F1 seeds were obtained by reciprocal crossing of WT with *arr16* mutant lines (*arr16 #1* and *arr16 #2*) following manual emasculation and pollination before flowering. T-DNA insertion mutants of *ARR16* in the Col-0 background (SALK_142105 and SALKseq_112294.2) were obtained from the NASC and homozygous lines were identified by PCR. The *ahk2/3, ahk2/4* and *ahk3/4* mutants (*ahk2-5/ahk3-7, ahk2-5/cre1-2* and *ahk3-7/cre1-2*, respectively) in Col-0 background were isolated as described previously^19^. All primers used for genotyping are listed in Supplemental Table 4.

### Plant growth conditions

Seeds were surface-sterilized and sown on Arabidopsis Gamborg B5 medium (Duchefa; http://www.duchefa.com) supplemented with 30 mM Suc. After stratification at 4°C in the dark for 3 days, plates were transferred to a growth chamber (18°C, 60% relative humidity, 16 h photoperiod, 50 μmol m^−2^ s^−1^ light intensity). For seed production, one-week-old seedlings were transplanted to non-enriched compost (Stender Substrates; https://www.stender.de/) in a glasshouse (18°-28°C) with a minimum photoperiod of 13 hours provided by supplementary lighting and plants were watered with Plan-Prod nutritive solution (Fertil; http://www.plantprod.com/).

### Analysis of seed coat permeability

Seed coat permeability was assessed by staining seeds with 1% (w/v) 2,3,5-triphenyltetrazolium chloride (Sigma-Aldrich; https://www.sigmaaldrich.com) solution at 30 °C in the dark for the time indicated in each figure. Formazan was extracted from the incubated seeds as described previously^31^ and the concentration was determined by measuring the absorbance at 485 nm with a SPECTROstar Nano microplate reader (BMG Labtech; https://www.bmglabtech.com). The RGB color image showing about 100 staining seeds was acquired by Axio Zoom.V16 microscope (Zeiss) and the stained area (%) of seeds was quantified with Fiji software^32^ (ImageJ version 1.52p / Java version 1.8.0_66) (Supplemental Fig. 1).

### Physiological analysis of seed traits

A thousand seeds were counted using the Elmor C3 seed counter (Elmor; https://elmor.com) and weighed on an electronic balance. The seed size was calculated by analyzing an image of about 100 seeds obtained using an Axio Zoom.V16 microscope with Fiji software. To estimate seed longevity, a controlled deterioration treatment (CDT) was performed as described previously^33^. Briefly, dry mature seeds were equilibrated at 80% relative humidity and 20°C for 3 days. The deterioration treatment was carried out at 80% relative humidity and 40°C for 0, 3, 5 and 7 days, followed by re-equilibration at 32% relative humidity and 20°C for 3 days. Germination assays were conducted using 0.8% (w/v) agarose plates with 50 seeds in a growth chamber at 18°C for 7 days, after 3 days stratification as described above. At 7 days, seed survival rates were scored based on the opening of healthy, green cotyledons following radicle emergence, since CDT often inhibits seedling establishment without affecting radicle emergence. The time for survival rate to decrease to 50% (P_50_) was estimated by the package viabilitymetrics^34^ (version 0.0.0.9100) in R^35^ (version 3.6.1). The speed of cotyledon greening (T_50CG_) was calculated based on the parameter t50.total with FourPHFfit.bulk function in the R package germinationmetrics^36^ (version 0.1.3.9000).

### Suberin analysis

Mature dry seeds were examined under UV illumination to visualize the autofluorescence (AF) in the hilum region of the seed as an indicator of suberin deposition^5^ using an Axio Zoom.V16 microscope. Sudan Red 7B lipid-staining for suberin of the seed coat was performed with 0.1% (w/v) Sudan Red 7B in a 1:1 (v/v) solution of polyethylene glycol 400:glycerol^15^, and stained seeds were observed with an Axio Zoom.V16 microscope. The AF in the hilum and Sudan Red 7B staining were quantified with Fiji software, and standardizing for each seed by dividing the total intensity at the hilum region by the area value of the seed. The seed suberin composition was analyzed using 25 mg of dry seeds (n=5 per genotype) as previously described^31^.

### GWAS and haplotype analysis

GWAS was performed with the GWA-Portal (https://gwas.gmi.oeaw.ac.at/)^37^ in which values of the stained area (%) of seeds at 72 hours after TZ staining and AF levels in the hilum region of dry seeds for 145 natural accessions were used as phenotypic data without transformation. The 1001 full sequence data set TAIR 9 was set as genotype data, followed by association analyses using the linear regression model. Manhattan and Miami plots were generated using the R package qqman^38^ (version 0.1.4) with SNPs having minor allele frequencies > 0.1. The SNPs significantly associated with phenotypic data variations were detected by the Benjamini–Hochberg method (FDR < 0.01). Linkage disequilibrium and haplotype block analysis in the *ARR16* genetic region were performed using the software Haploview (version 4.2) with the confidence intervals method^39^ for 17 SNPs having minor allele frequencies > 0.1. Graphic representation of gene structural information were generated with Exon-Intron Graphic Maker (http://wormweb.org/exonintron). RNA secondary structure was visualized with ViennaRNA web services/forna (http://rna.tbi.univie.ac.at/forna/)^40^ and effects of SNPs on local RNA structure were predicted by RNAsnp web server (https://rth.dk/resources/rnasnp/)^41^.

### Quantitative RT-PCR

Developing seeds were freshly harvested and frozen in liquid nitrogen. About 40 mg of sample were ground to a powder with a mortar and pestle and mixed with an equal amount of polyvinylpolypyrrolidone (Sigma-Aldrich), then total RNA was extracted using Rneasy Plant Mini Kit (Qiagen; https://www.qiagen.com), according to the manufacturer’s protocol including the options to add 20 μl of 2 M dithiothreitol (Life Technologies/Thermo Scientific; https://www.thermofisher.com) per 1 ml buffer RLC and on-column Dnase digestion with Dnase I (Qiagen). First-strand cDNA was synthesized from 1 μg of total RNA using a Thermo Scientific RevertAid RT Kit (Thermo Scientific). Quantitative RT-PCR was performed using a CFX Connect Real-Time PCR Detection System (Bio-Rad; https://www.bio-rad.com) and SsoAdvanced PreAmp Supermix Biorad kit (Bio-Rad), according to the manufacturer’s protocol. The thermal cycling conditions were: 95°C for 8 min followed by 40 cycles of 10 s at 95°C for denaturation, and 10 s at 60°C for annealing and extension. The primers used for RT-qPCR were designed from the target gene sequences using Primer3 software (http://frodo.wi.mit.edu) and are listed in Supplementary Table 5. A constitutive gene *AT5G60390 (EF1α*) and *AT4G12590* that is stably expressed in seed^42^ were used as reference genes to calculate relative mRNA levels for all the samples.

### Data plots, statistical analysis and microarray database search

Bar graphs with jitter points were drawn using the package ggplot2^43^ (version 3.3.5) in R. Box plots and split violin plots with jitter points were made using the package Plotly^44^ (version 4.9.1) in R. Significant differences among three or more groups were evaluated by Tukey–Kramer test or Dunnett’s test. All statistical tests were performed using R. Expression profile of *ARR16* gene in developing seed tissues based on the microarray analysis^45^ was obtained from the Arabidopsis eFP Browser^46^ (https://bar.utoronto.ca/efp_arabidopsis/cgi-bin/efpWeb.cgi).

## Acknowledgments

We thank D. C. Bergmann (Stanford University) for the gift of *arr16#1* and *arr16#2* seeds, and T. Schmülling (Freie Universität Berlin) for the gift of *ahk2/3, ahk2/4* and *ahk3/4* seeds. We are also grateful to A. Berger, C. Sallé, A. Frey, A. To and I. Debeaujon in our institute for their technical aid and advice. N.S. received funding from the Japanese Society for the Promotion of Science (JSPS) Overseas Research Fellowship and the European Union’s Horizon 2020 research and innovation programme under the Marie Skłodowska-Curie grant agreement No [846387]. The IJPB benefits from the support of Saclay Plant Sciences-SPS (ANR-17-EUR-0007). The authors thank the Bordeaux-Metabolome Facility, supported by MetaboHUB (ANR-11-INBS-0010), where GC-based suberin analyses were performed.

