## Supplemental files for "Cytokinin response regulator *ARR16* regulates seed coat permeability in Arabidopsis natural accessions"

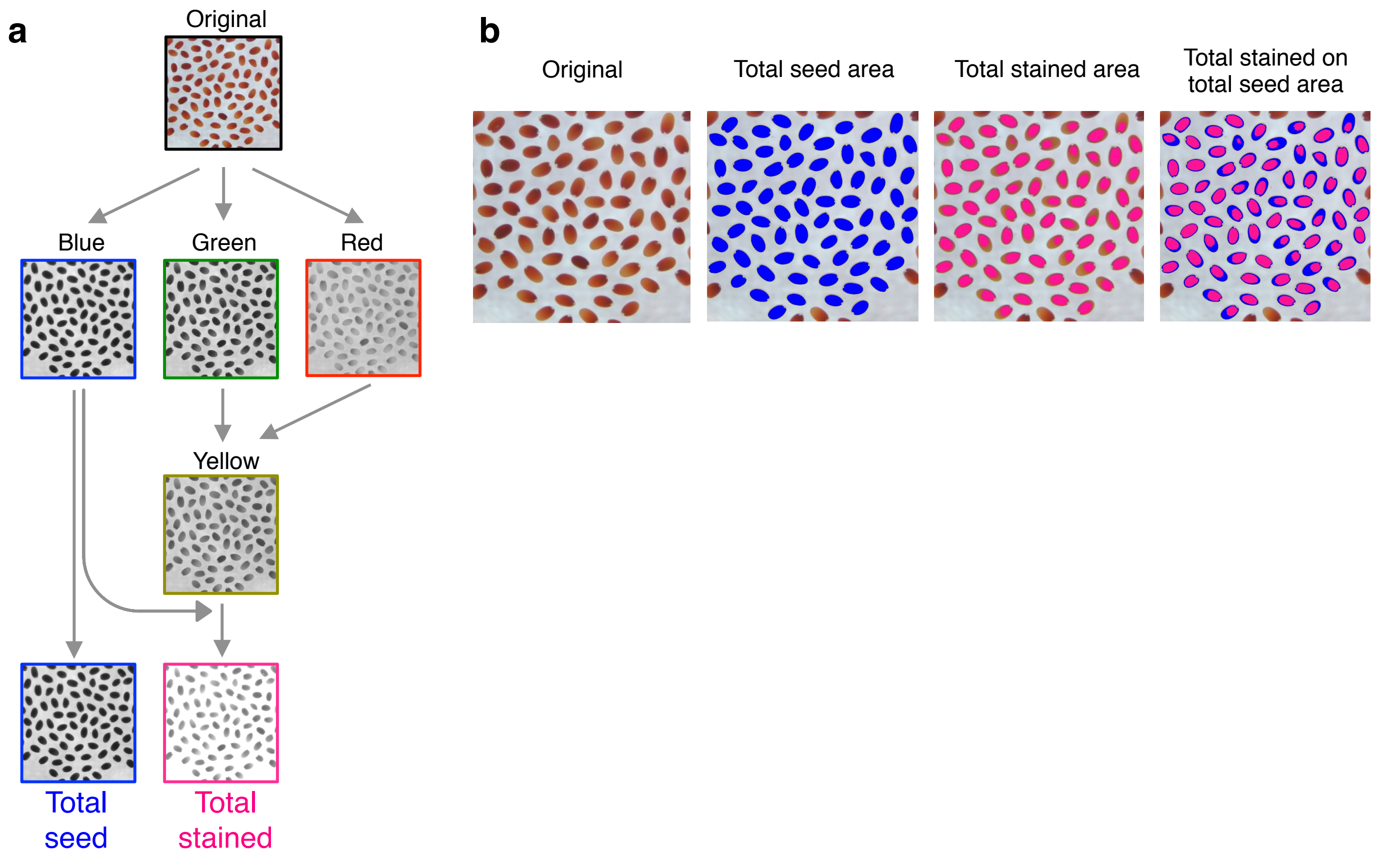


**Supplementary Figure 1. Quantitative method for determining seed coat permeability by image analysis using Fiji software. a**, Diagram of the image analysis procedure. Seeds are stained with 1% (w/v) aqueous tetrazolium (TZ) solution at 30°C in the dark. Original images (RGB, 24-bit, 300 DPI) of approximately 100 seeds were captured and stored. The original image was split into red, green and blue channels. The blue channel was used for measurement of total seed area because the TZ stained seeds have little blue color; this is converted to black. The red and green images were averaged together to produce a yellow image that corresponds to the natural unstained color of the seeds. The yellow image was then added to blue image in order to extract stained area (non-yellow like area) as a black and white image by using Image Calculator function. The black-and-white regions were then inverted before binarization. **b**, Overlays of binarized images compared to the original image. Binarization was performed using the "Threshold" command and either the method "Li" for total seed area or the method "Default: 80* to 255" for total stained area, respectively. Regions of interest on binarized images were detected by the "Analyze Particles" command and resulting regions are shown as overlays on the original image. Blue and pink regions indicate total seed area and total stained area, respectively. The stained rate as an index for seed coat permeability was calculated using the formula; Stained area (%) = (total stained area / total seed area) × 100.

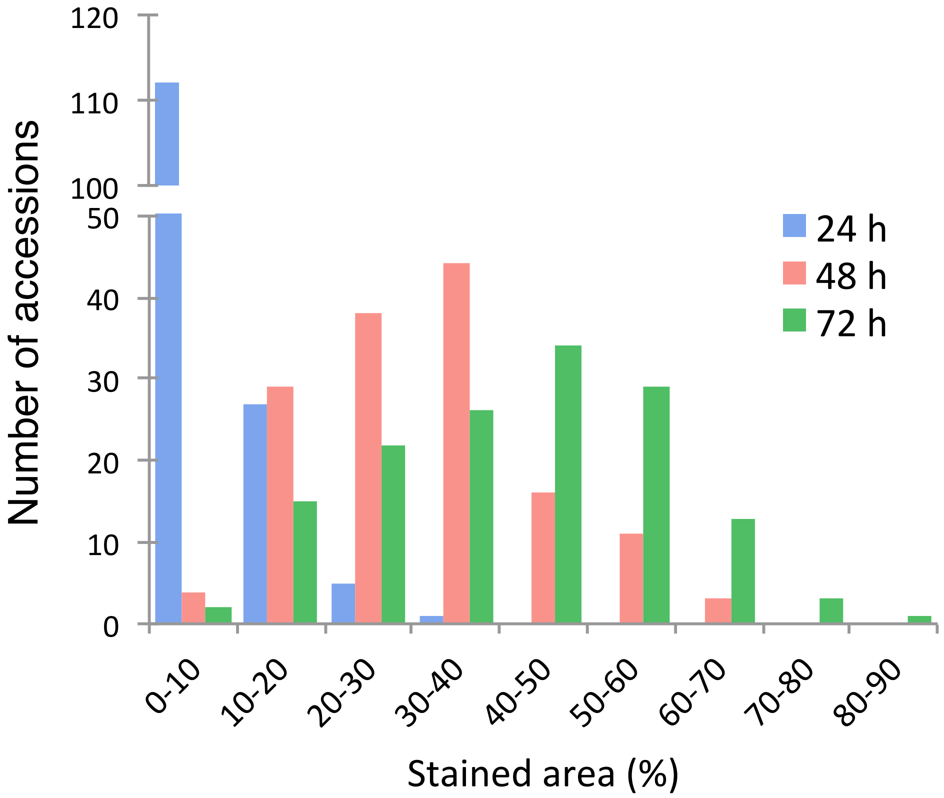


**Supplementary Figure 2. Frequency distribution of stained area (%) for 145 natural accession seeds stained with TZ solution.** Seeds were stained for 24, 48 and 72 hours and stained area (%) was quantified by image analysis.

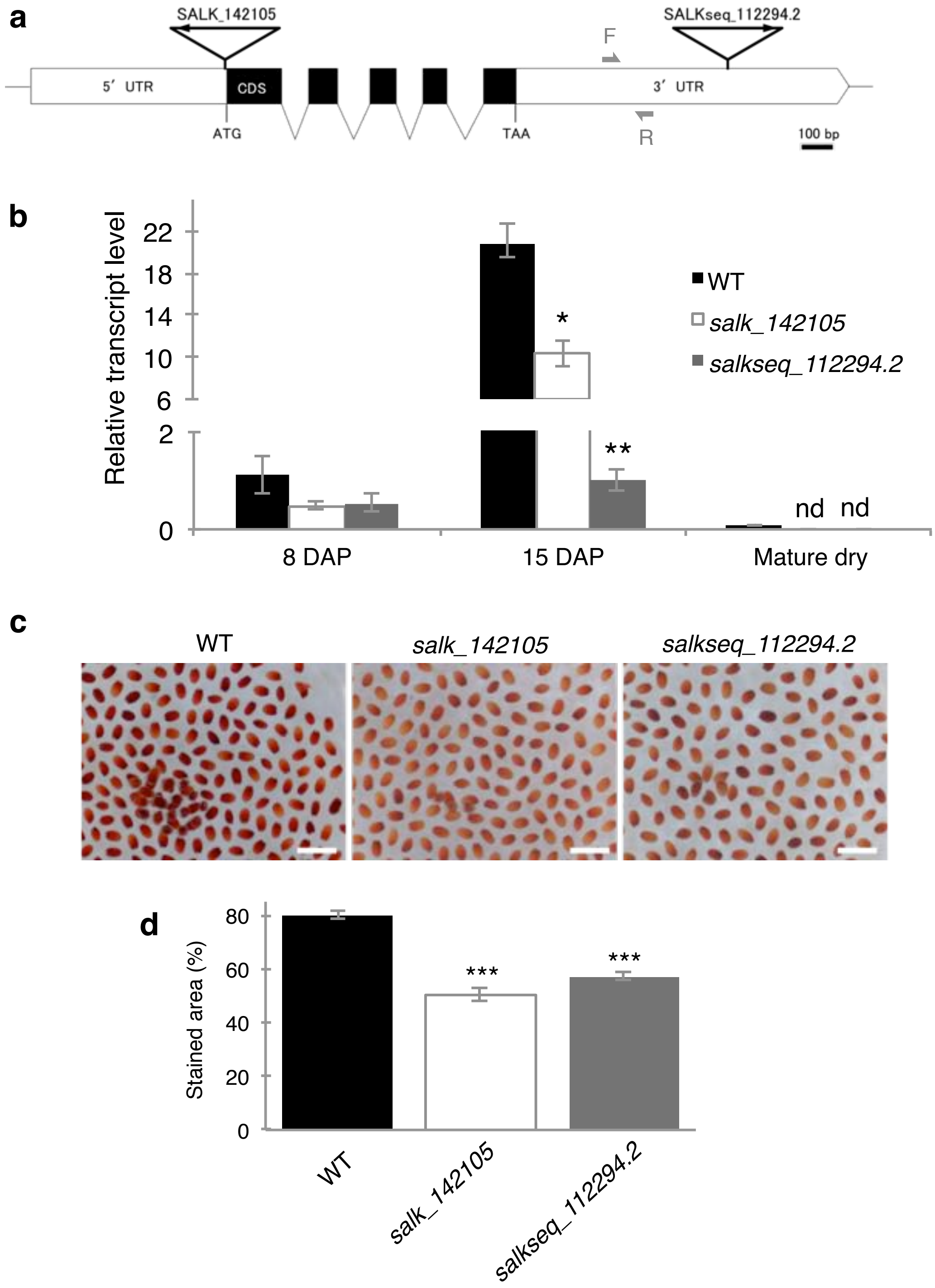


**Supplementary Figure 3. Effect of T-DNA insertions in *ARR16* gene on gene expression and seed coat permeability.** **a**, Schematic representation of the position of T-DNA insertions in the *ARR16* gene. ATG; start codon, TAA; stop codon, UTR; untranslated region, CDS; coding sequence, diagonal lines; introns. T-DNA insertions are indicated with black triangles. Gray arrows F and R show position of forward and reverse primers, respectively, used for RT-qPCR analysis. Scale bars represent 100 base pairs. The graphic was generated with the Exon-Intron Graphic Maker (http://wormweb.org/exonintron). **b**, *ARR16* expression in WT and T-DNA insertion lines (*salk_142105* and *salkseq_112294.2*) during seed development at 8, 15 days after pollination (DAP) and in mature dry seeds analyzed by RT-qPCR. Values presented are means ±SE (n=3) (*$P < 0.05$, **$P < 0.01$, Dunnett's test). nd; not detected. **c**, Seeds of WT and T-DNA insertion lines at 72 hours after TZ staining. Scale bars represent 1 mm. **d**, Stained area (%) of WT and T-DNA insertion line seeds at 72 hours after TZ staining. Values presented are means ±SE (n=3) (***$P < 0.001$, Dunnett's test).

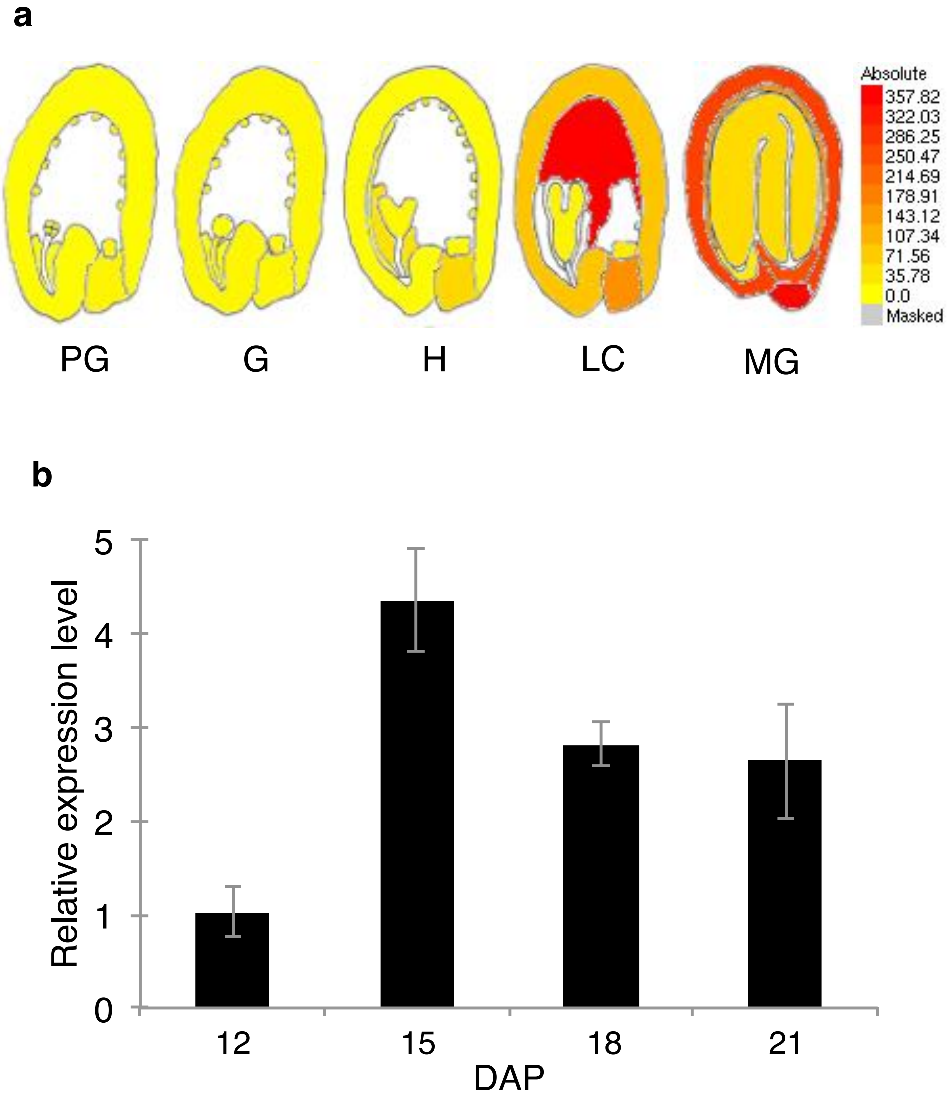


**Supplementary Figure 4. *ARR16* gene expression during seed development. a**, Expression profile of *ARR16* gene in developing seed tissues obtained from The Arabidopsis eFP Browser (Winter et al. 2007 using data from Le et al. 2010; https://bar.utoronto.ca/efp_arabidopsis/cgi-bin/efpWeb.cgi). PG; pre-globular, G; globular, H; heart, LC; linear cotyledon and MG; mature green fruit stages of seed development. **b**, Steady-state levels of *ARR16* transcripts during seed development at 12, 15, 18 and 21 days after pollination (DAP) analyzed by RT-qPCR. Values presented are means ±SE (n=3).

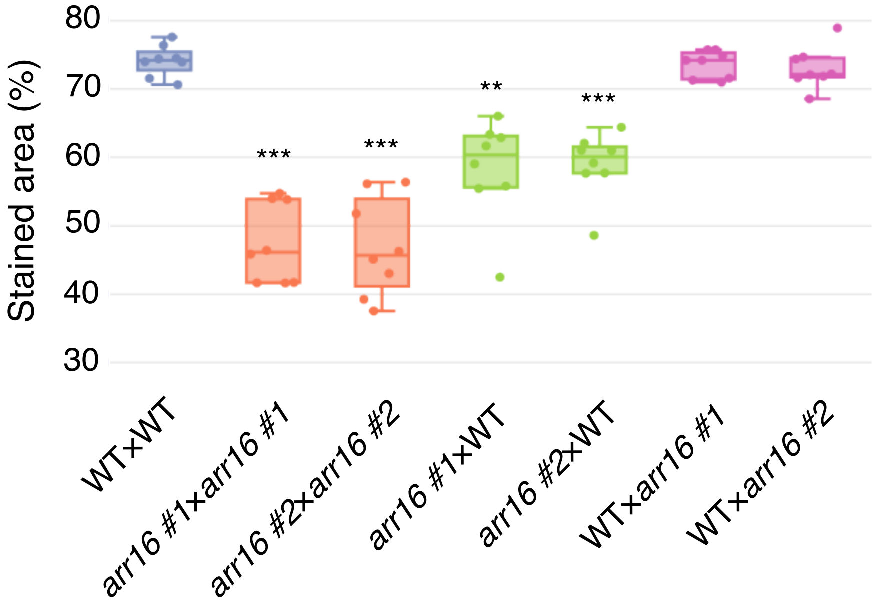


**Supplementary Figure 5. Maternal effect of reduced seed coat permeability in *arr16*.** $F_1$ seeds were obtained by reciprocal crossing of WT and *arr16* mutant lines following manual emasculation and pollination before flowering. Seeds from WT and *arr16* mutants, and $F_1$ seeds from reciprocal crosses were subjected to TZ staining for 72 hours. $F_1$ seeds are denoted by the maternal genotype followed by the paternal genotype. Boxplot with jitter points ($n$ = 8) showing the stained area (%) of seeds quantified by image analysis (**$P < 0.01$, ***$P < 0.001$, Dunnett's test, compared to WT(♀)×WT(♂)).

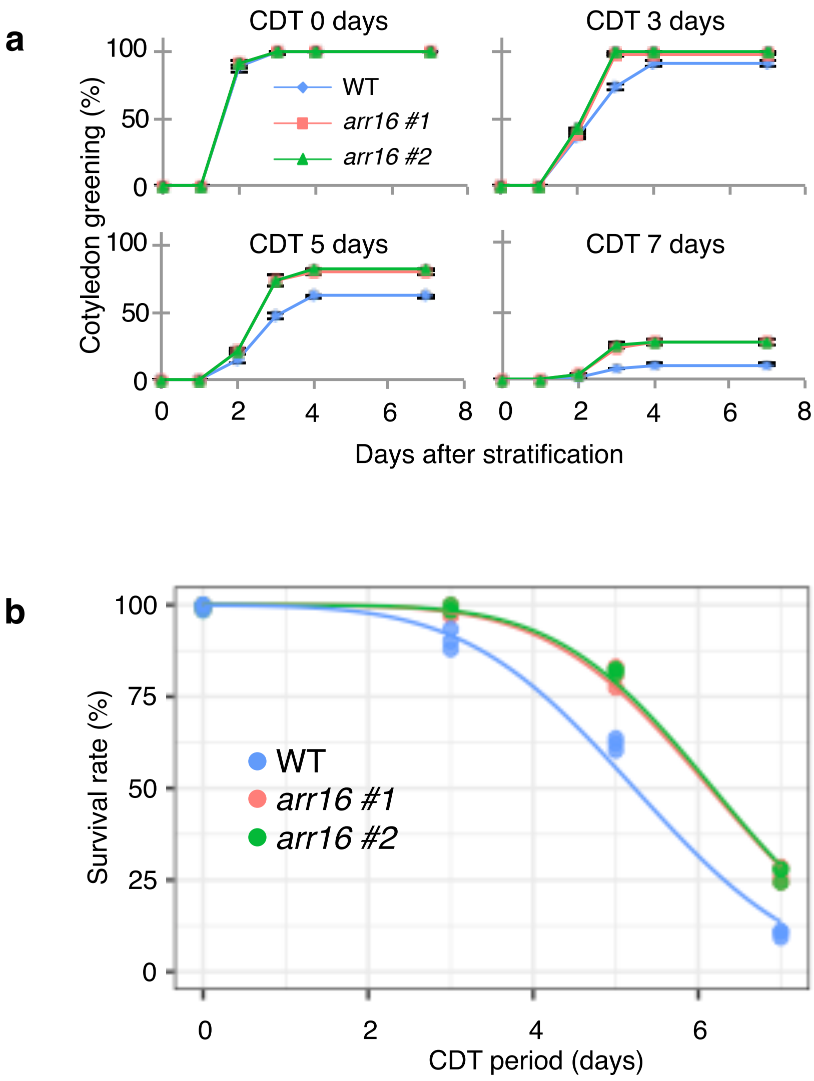


**Supplementary Figure 6. Seed vigor and longevity of *arr16* after CDT. a,** Establishment of seedlings of WT and *arr16* mutant seeds after the controlled deterioration treatment (CDT). CDT was performed for 0, 3, 5 and 7 days, followed by stratification and germination tests. Healthy seedling establishment after germination was scored based on the opening of green cotyledons following radicle emergence, since CDT often inhibits seedling establishment without affecting radicle emergence. Values presented are means ±SE (n=3). **b,** Survival curves of WT and *arr16* mutant seeds following CDT. Seed survival rates were scored based on the opening of green healthy cotyledons after 7 days in the germination assays presented in (a).

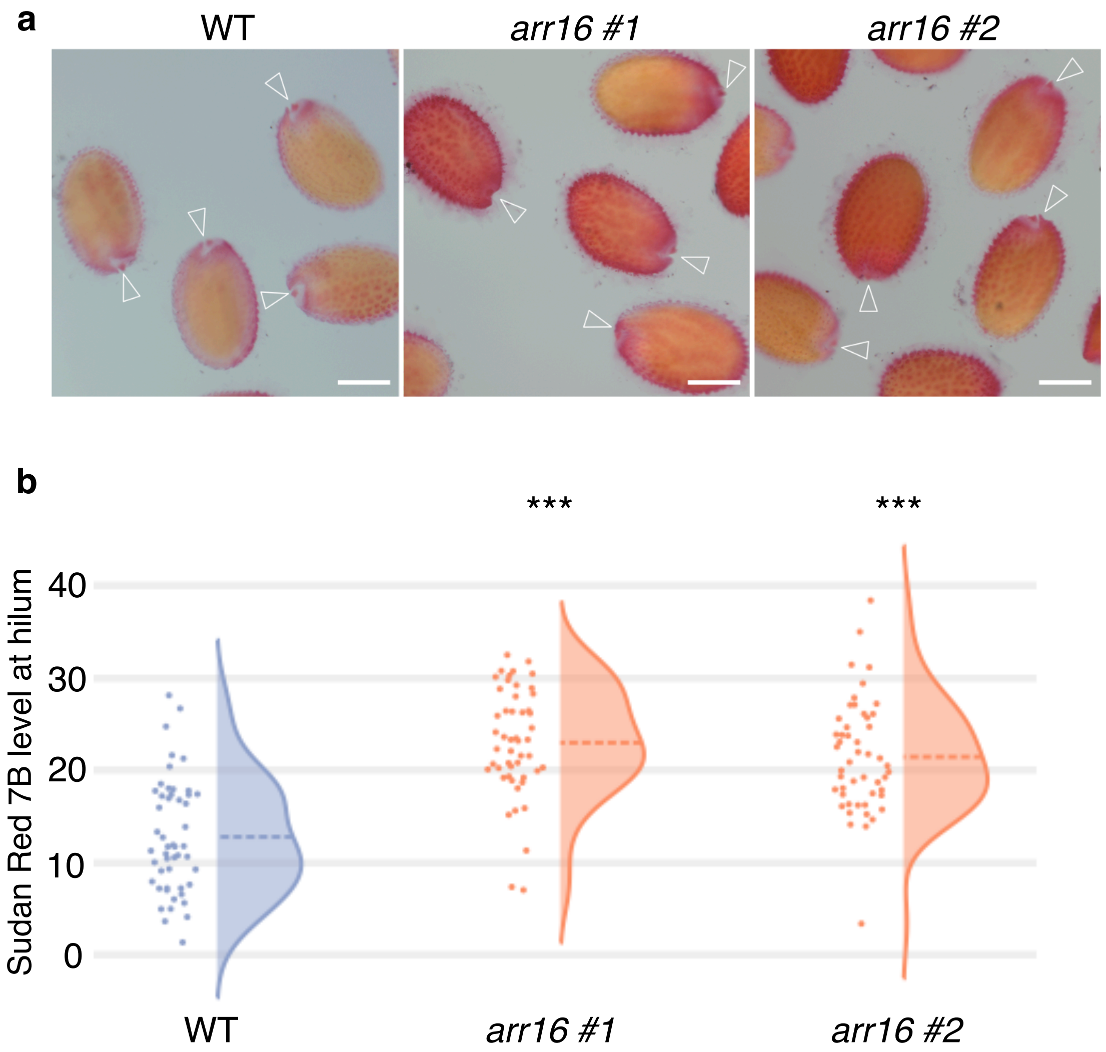


**Supplementary Figure 7. Staining of *arr16* seeds with the lipophilic dye Sudan Red 7B. a,** Seeds of WT and *arr16* mutants at 24 hours after Sudan Red 7B staining. White arrowhead indicates the position of the hilum. Scale bars represent 200 µm. **b**, Split violin plots with rotated kernel density estimates (dashed line represents mean) and jitter points (n=50) showing the staining level in hilum region of WT and *arr16* mutant seeds at 24 hours after Sudan Red 7B staining (***$P < 0.001$, Dunnett's test).

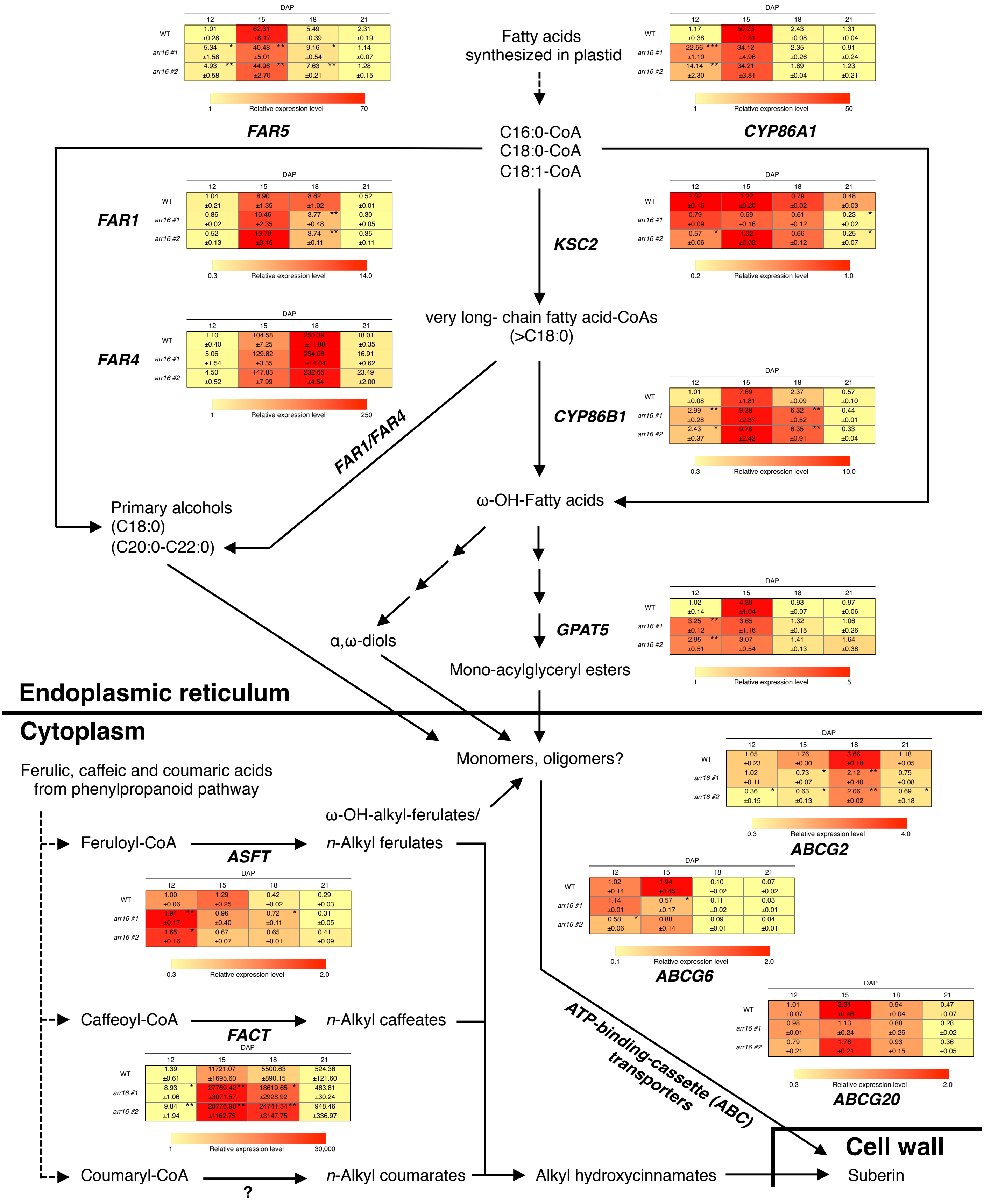


**Supplementary Figure 8. Effect of *ARR16* mutation on the expression of genes encoding proteins involved in suberin biosynthesis.** Transcript expression levels in developing seeds of WT and *arr16* mutants at 12, 15, 18 and 21 days after pollination (DAP) were analyzed by RT-qPCR. Earlier induction of gene expression was observed in *arr16* compared to WT for genes such as *FAR5*, *CYP86A1*, *CYP86B1*, *GPAT5*, *ASFT* and *FACT* at 12 DAP. *CYP86B1* and *FACT* transcript levels were also significantly higher in *arr16* later at 18 DAP, while the transcript abundance of ABCG2/6/20 transporters, which act to promote suberin deposition, was slightly lower in *arr16* than WT. Values presented in tables are means ±SE (n=3) (**P* < 0.05, ***P* < 0.01, ****P* < 0.01, Dunnett's test). The overview of the suberin biosynthetic pathway was modified from Vishwanath *et al.* *Plant Cell Rep* 34, 573–586 (2015).

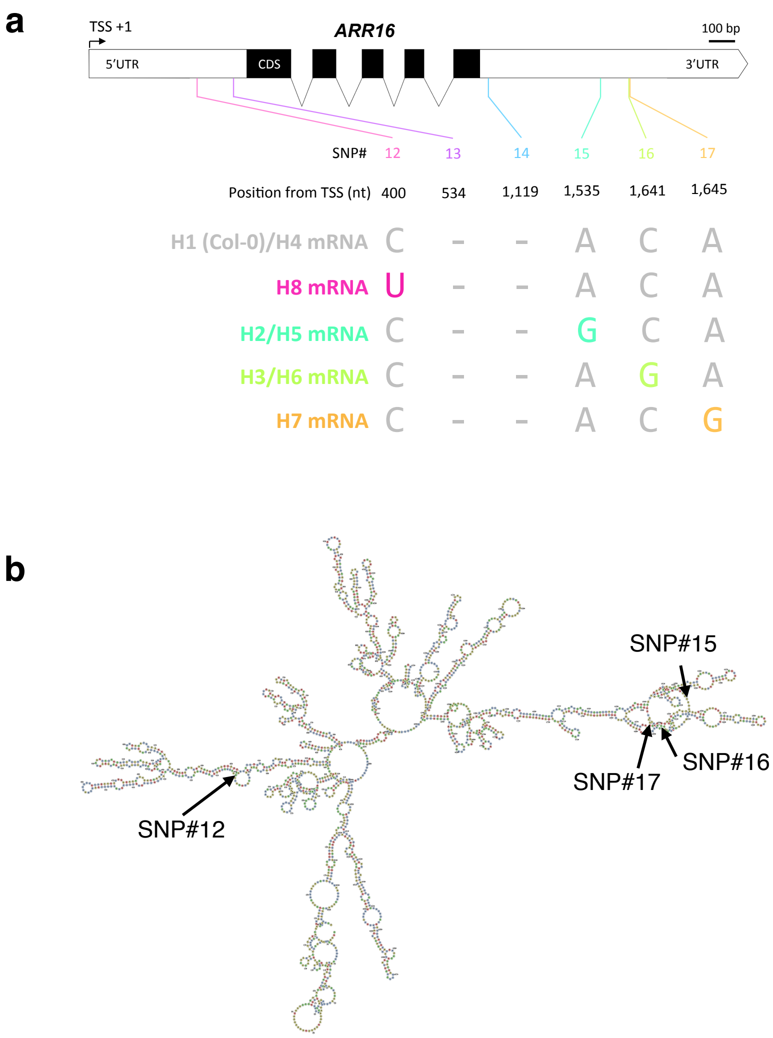


**Supplementary Figure 9. SNPs in the UTRs of *ARR16* gene and the SNP positions in the mRNA secondary structure. a**, Schematic representation of the SNPs position in the UTRs of *ARR16* and the corresponding haplotypes. TSS; transcription start site, UTR; untranslated region, CDS; coding sequence, diagonal lines; introns. nt; nucleotide. Scale bars represent 100 base pairs. The graphic was generated using Exon-Intron Graphic Maker (http://wormweb.org/exonintron). The *ARR16* gene, including the promoter region, has eight haplotypes (Fig. 4b), and these haplotypes were classified into five groups (H1/H4, H2/H5, H3/H6, H7 and H8) based on the SNPs for mRNA sequence, which excludes the promoter region. **b**, Predicted secondary structure of *ARR16* mRNA. The full-length mRNA sequence (5′UTR +CDS +3′UTR; 2,073 nt) from the Col-0 accession, a H1 group member, was used for the prediction. The black arrows indicate the positions of SNPs.

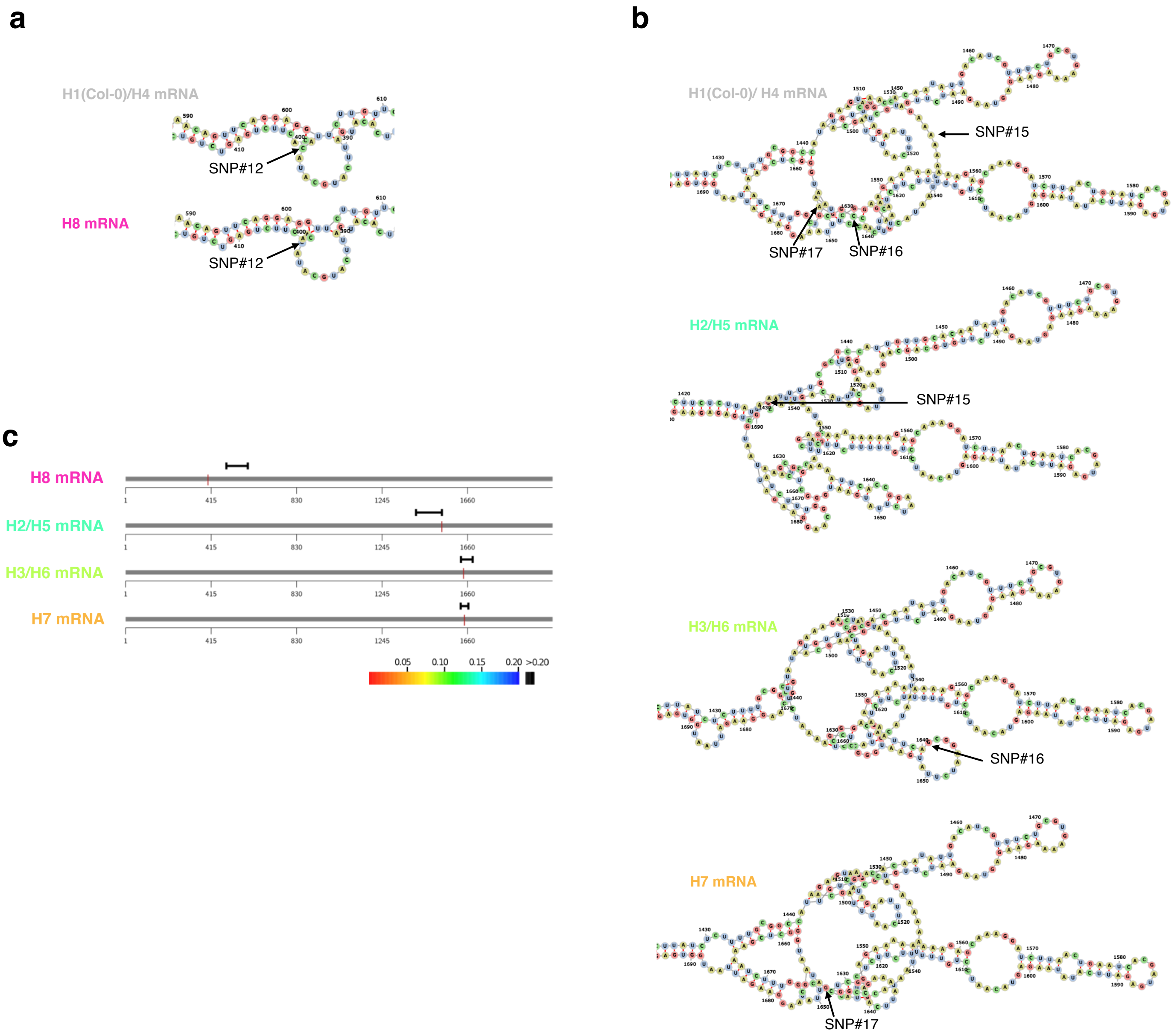


**Supplementary Figure 10. Predicting the effect of natural variant SNPs on the secondary structure of *ARR16* mRNA. a**, Comparison of predicted RNA structures neighboring SNP#12 (H1/H4 vs H8). **b**, Comparison of predicted RNA structures neighboring SNP#15 (H1/H4 vs H2/H5), SNP#16 ((H1/H4 vs H3/H6) and SNP#17 (H1/H4 vs H7). **c**, Statistical analysis of *ARR16* mRNA structural changes caused by natural variant SNPs. Each haplotype mRNA was compared to a reference H1/H4 mRNA sequence (2,073 nt). Red vertical line indicates the position of the SNP. The local region detected with maximum structural change is colored according to the *P*-value according to the rainbow scale. If the *P*-value is greater than 0.2, the region is colored in black (*i.e.* no significant structural change).

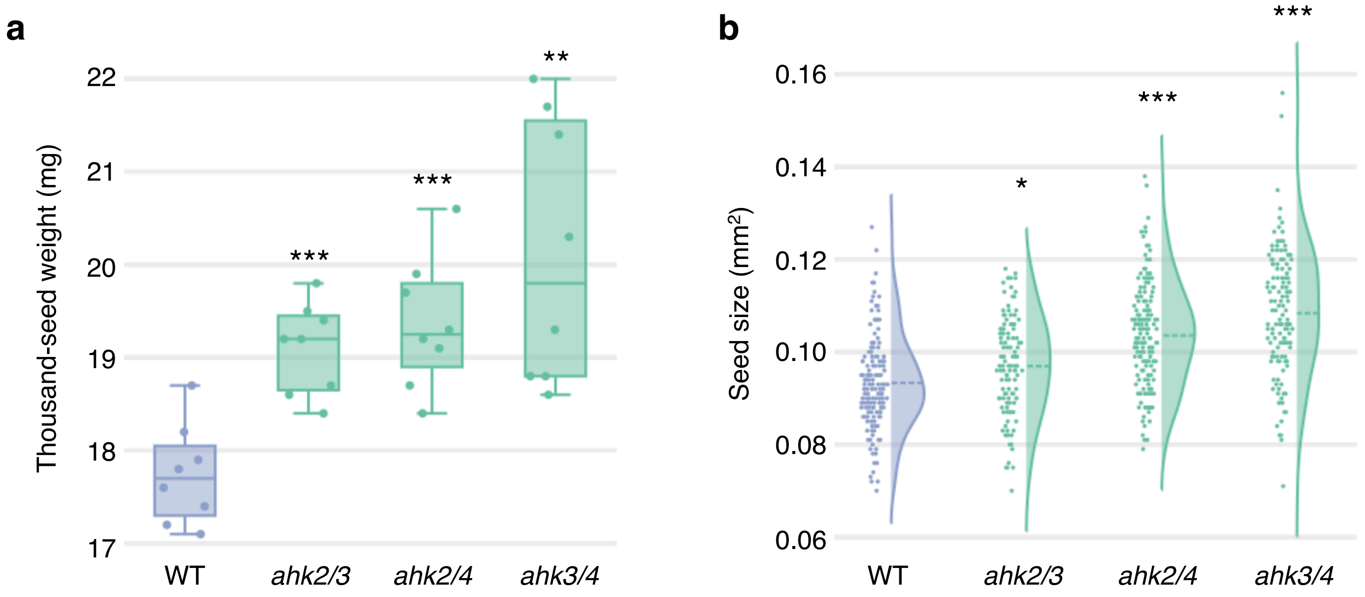


**Supplementary Figure 11. Seed weight and size of *ahk* double mutants. a**, Boxplot with jitter points (n=8) showing thousand-seed weight of WT, *ahk2/3*, *ahk2/4* and *ahk3/4* seeds (***P* < 0.01, ****P* < 0.001, Dunnett's test). **b**, Split violin plots with rotated kernel density estimates showing seed size of WT, *ahk2/3*, *ahk2/4* and *ahk3/4* (**P* < 0.05, ****P* < 0.001, Dunnett's test). Jitter points represent seed size data (*n* = 143, 113, 153 and 128 for WT, *ahk2/3*, *ahk2/4* and *ahk3/4*, respectively). Dashed line shows the mean value for each genotype.

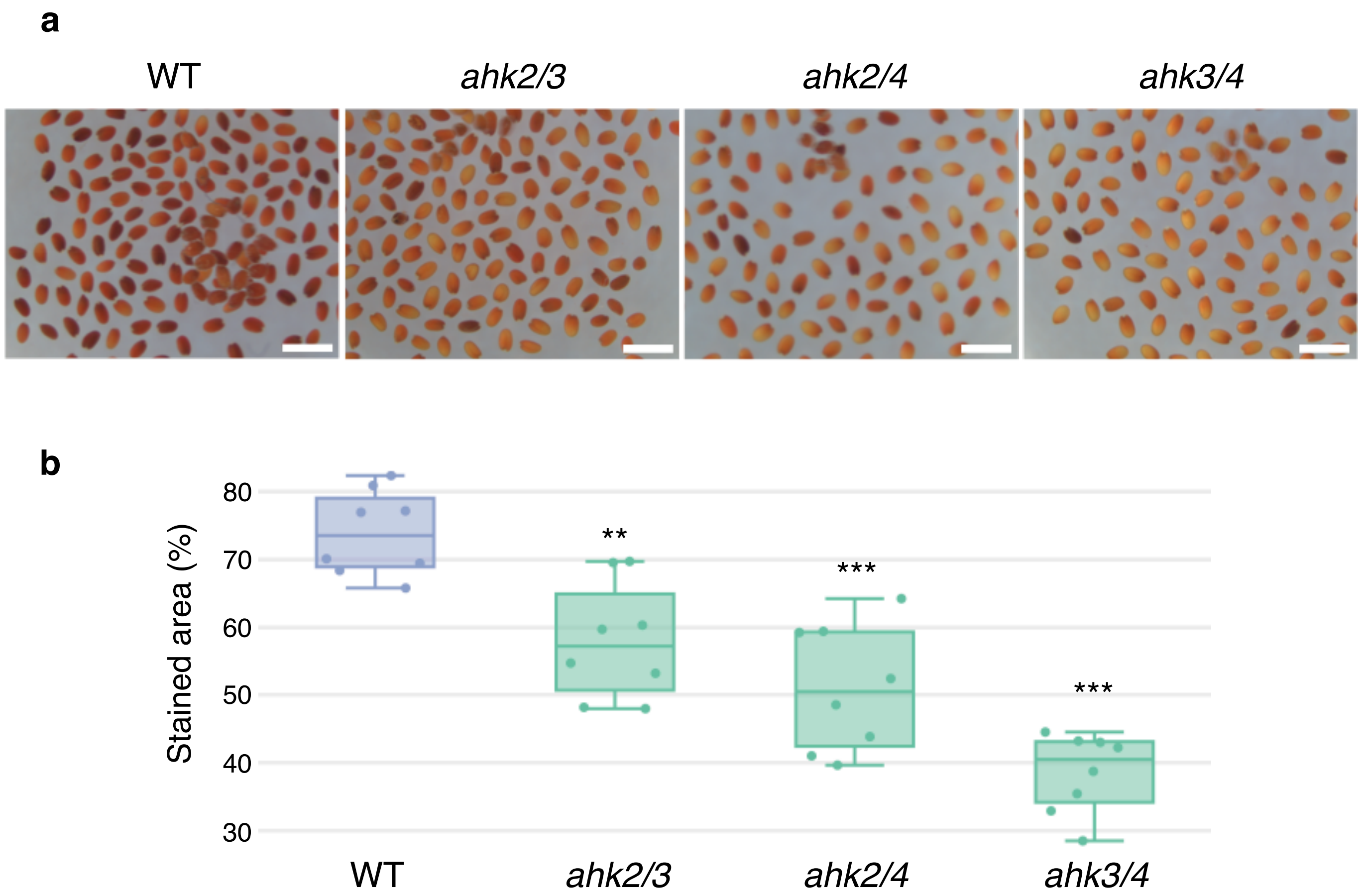


**Supplementary Figure 12. Seed coat permeability of *ahk* double mutants. a**, Seeds of WT, *ahk2/3*, *ahk2/4* and *ahk3/4* at 72 hours after tetrazolium (TZ) staining. Scale bars represent 1 mm. **b**, Boxplot with jitter points (n=8) showing stained area (%) of WT, *ahk2/3*, *ahk2/4* and *ahk3/4* seeds at 72 hours after TZ staining (**$P < 0.01$, ***$P < 0.001$, Dunnett's test).

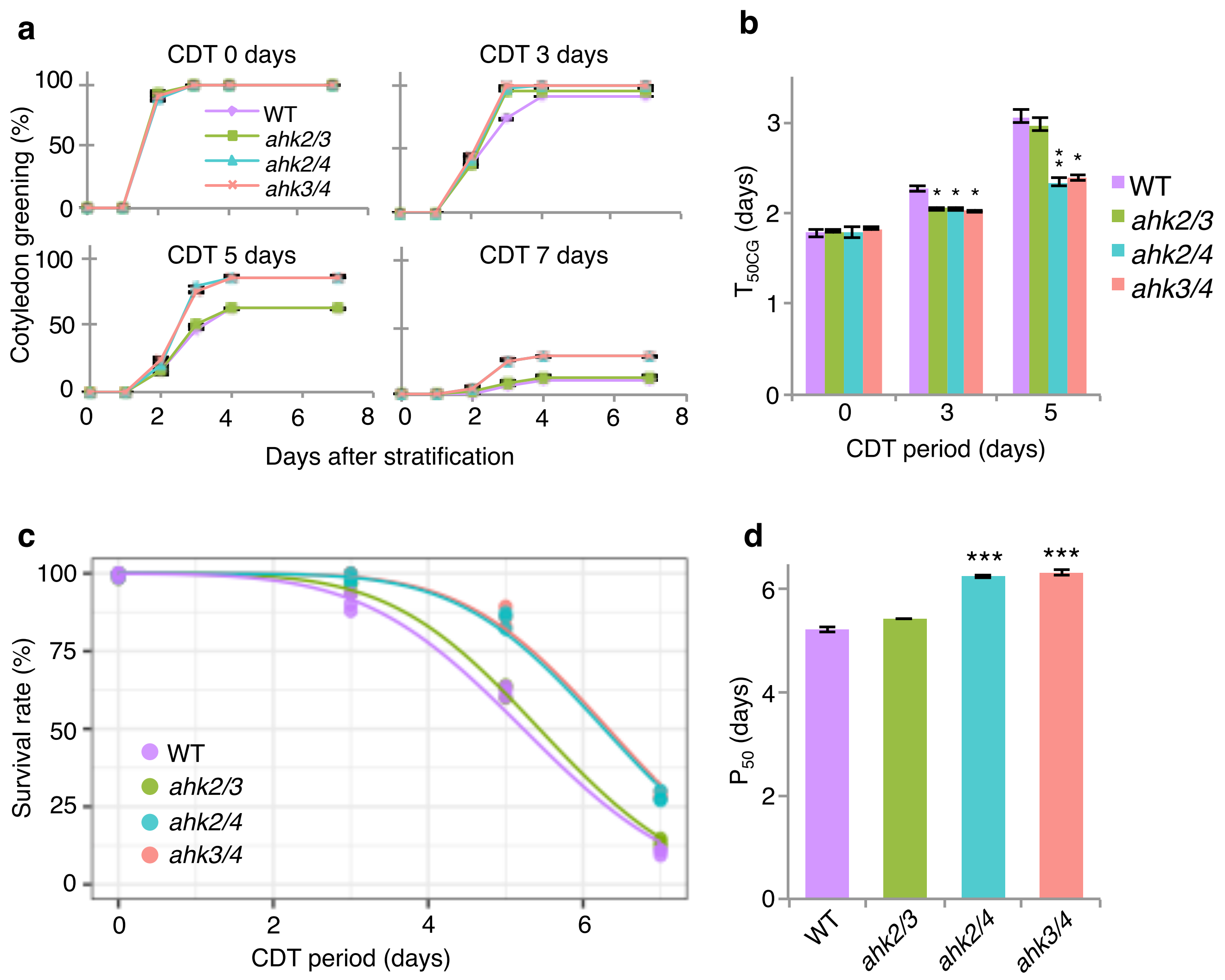


**Supplementary Figure 13. Seed vigor and longevity of *ahk* double mutants after CDT. a**, Establishment of seedlings of WT and *ahk* double mutant seeds after a controlled deterioration treatment (CDT). CDT was performed for 0, 3, 5 and 7 days, followed by stratification and germination tests. Healthy seedling establishment after germination was scored based on the opening of green cotyledons following radicle emergence, since CDT often inhibits seedling establishment without affecting radicle emergence. **b**, Cotyledon greening speed ($T_{50CG}$; time required for 50% of viable seeds to develop green cotyledons) at 0, 3 and 5 days after CDT. Values presented are means ±SE (n=3) (*$P < 0.05$, **$P < 0.01$, Dunnett's test). **c**, Survival curves of WT and *ahk* double mutant seeds in CDT. Seed survival rates were scored based on the opening of healthy, green cotyledons after 7 days in the germination assays in (a). **d**, Index of seed longevity $P_{50}$ ; time for survival rate to decrease by 50%. Values presented are means ±SE (n=3) (***$P < 0.001$, Dunnett's test).

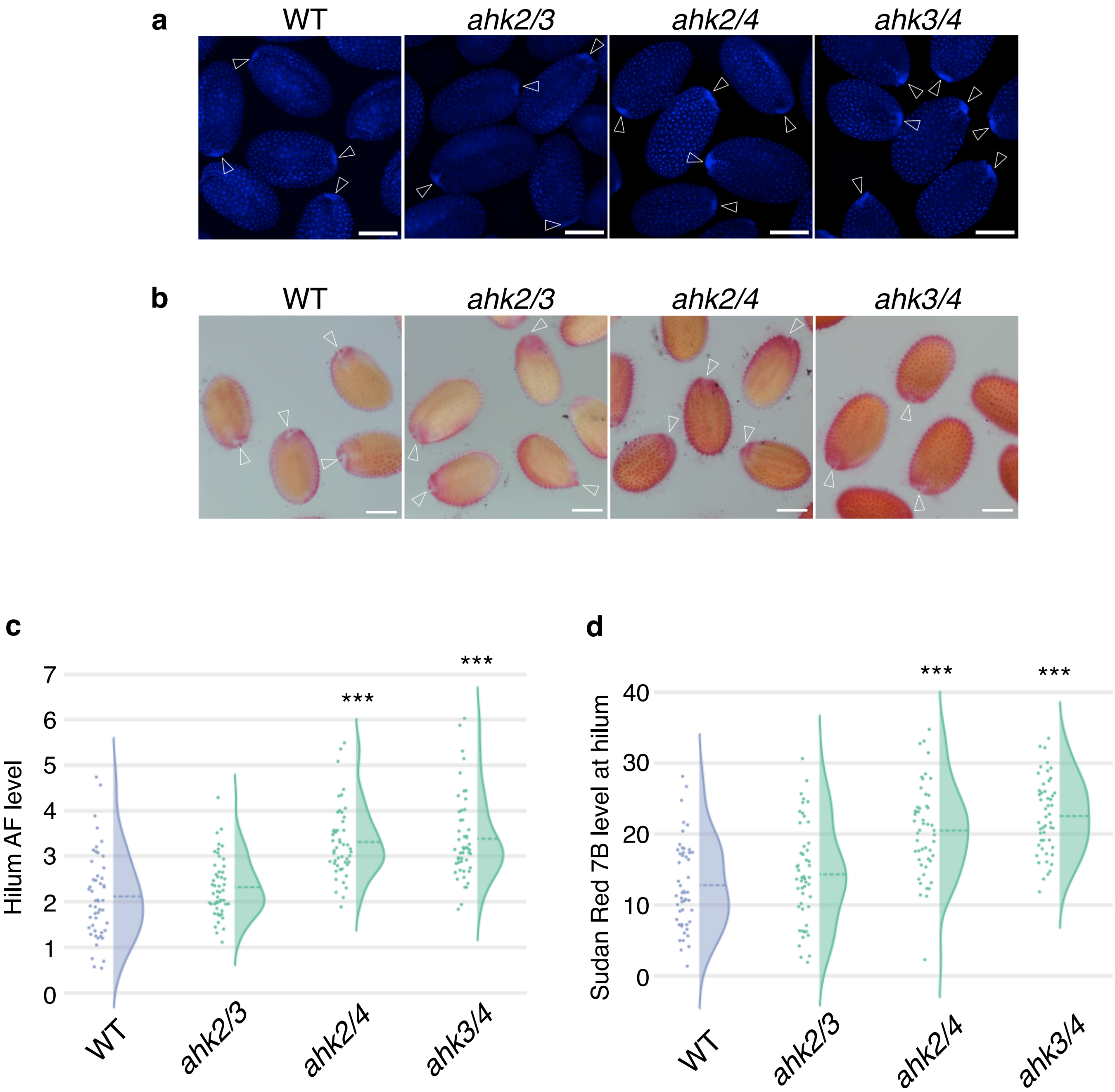


**Supplementary Figure 14. Suberin accumulation in the hilum region of *ahk* mutant seeds.** Dry seeds under UV light (**a**) and seeds stained with Sudan Red 7B at 24 hours after staining (**b**) in WT, *ahk2/3*, *ahk2/4* and *ahk3/4*. White arrowhead indicates the position of the hilum. Scale bars represent 200 µm. Split violin plots with rotated kernel density estimates (dashed line represents mean) and jitter points (n=50) showing the autofluorescence (AF) level in the hilum region of dry seeds (**c**) and the staining level in the hilum region 24 hours after Sudan Red 7B staining (**d**) in WT, *ahk2/3*, *ahk2/4* and *ahk3/4* (***$P <$ 0.001, Dunnett's test). Dashed line shows the mean value for each genotype.

**Supplementary Table 1.** 145 natural accessions and phenotype data used in GWAS for seed coat permeability. Accessions information was obtained from the 1001 Genomes (https://1001genomes.org/).
Country of origin is indicated by ISO 3166 code (https://www.iso.org/iso-3166-country-codes.html).

| Accession ID | Name | Country of origin | Stained area (%) at 72 h after TZ staining |
|---|---|---|---|
| 430 | Gr-1 | AUT | 50.04 |
| 763 | Kar-1 | KGZ | 60.98 |
| 765 | Sus-1 | KGZ | 50.26 |
| 766 | Dja-1 | KGZ | 53.64 |
| 768 | Zal-1 | KGZ | 43.48 |
| 772 | Neo-6 | TJK | 60.93 |
| 5837 | Bor-1 | CZE | 34.18 |
| 6897 | Ag-0 | FRA | 54.52 |
| 6898 | An-1 | BEL | 19.29 |
| 6903 | Bor-4 | CZE | 53.18 |
| 6908 | CIBC-5 | UK | 26.27 |
| 6909 | Col-0 | USA | 69.83 |
| 6915 | Ei-2 | GER | 39.51 |
| 6919 | Ga-0 | GER | 53.61 |
| 6922 | Gu-0 | GER | 41.06 |
| 6924 | HR-5 | UK | 36.21 |
| 6926 | Kin-0 | USA | 39.68 |
| 6929 | Kondara | TJK | 55.49 |
| 6931 | Kz-9 | KAZ | 57.61 |
| 6940 | Mz-0 | GER | 28.55 |
| 6945 | Nok-3 | NED | 15.40 |
| 6951 | Pu2-23 | CZE | 13.02 |
| 6956 | Pu2-7 | CZE | 57.49 |
| 6958 | Ra-0 | FRA | 51.26 |
| 6959 | Rennes-1 | FRA | 49.28 |
| 6961 | Se-0 | ESP | 43.05 |
| 6963 | Sorbo | TJK | 65.69 |
| 6967 | Sq-8 | UK | 52.08 |
| 6970 | Ts-1 | ESP | 32.31 |
| 6975 | Uod-1 | AUT | 40.58 |
| 6979 | Wei-0 | SUI | 61.20 |
| 6981 | Ws-2 | RUS | 8.51 |
| 6982 | Wt-5 | GER | 49.89 |
| 6984 | Zdr-1 | CZE | 39.52 |
| 6986 | Abd-0 | UK | 41.49 |
| 6987 | Ak-1 | GER | 61.62 |
| 6989 | Alst-1 | UK | 41.20 |
| 6990 | Amel-1 | NED | 38.75 |
| 6992 | Ang-0 | BEL | 19.03 |
| 6997 | Appt-1 | NED | 13.16 |
| 7000 | Aa-0 | GER | 52.23 |
| 7002 | Baa-1 | NED | 58.13 |
| 7003 | Bs-1 | SUI | 41.55 |
| 7013 | Bd-0 | GER | 60.67 |
| 7025 | Bl-1 | ITA | 45.61 |
| 7026 | Boot-1 | UK | 44.52 |
| 7028 | Bch-1 | GER | 23.09 |
| 7031 | Bsch-0 | GER | 30.71 |
| 7061 | Cal-0 | UK | 82.52 |
| 7062 | Ca-0 | GER | 45.19 |
| 7064 | Cnt-1 | UK | 65.52 |
| 7071 | Chat-1 | FRA | 37.48 |
| 7072 | Chi-0 | RUS | 52.27 |
| 7092 | Com-1 | FRA | 38.51 |
| 7096 | Di-G | FRA | 26.34 |
| 7103 | Dra-0 | CZE | 48.52 |
| 7106 | Dr-0 | GER | 40.52 |
| 7107 | Durh-1 | UK | 37.75 |
| 7109 | Ema-1 | UK | 14.52 |
| 7117 | El-0 | GER | 16.18 |
| 7119 | En-2 | GER | 14.70 |
| 7120 | En-D | GER | 20.20 |
| 7125 | Er-0 | GER | 49.84 |
| 7126 | Es-0 | FIN | 62.46 |
| 7127 | Est | EST | 55.87 |
| 7133 | Fr-2 | GER | 32.05 |
| 7138 | Fi-0 | GER | 47.18 |
| 7143 | Gel-1 | NED | 17.58 |
| 7147 | Gie-0 | GER | 62.52 |
| 7161 | Gd-1 | GER | 29.98 |

**Supplementary Table 1.** Continued.

| Accession ID | Name | Country of origin | Stained area (%) at 72 h after TZ staining |
|---|---|---|---|
| 7162 | Hs-0 | GER | 20.89 |
| 7163 | Ha-0 | GER | 29.94 |
| 7165 | Hn-0 | GER | 55.65 |
| 7177 | Jm-0 | CZE | 25.08 |
| 7192 | Kil-0 | UK | 39.30 |
| 7199 | Kl-5 | GER | 47.74 |
| 7202 | Kb-0 | GER | 31.38 |
| 7203 | Krot-0 | GER | 24.03 |
| 7207 | Kyoto | JPN | 31.14 |
| 7208 | Lan-0 | UK | 47.78 |
| 7209 | La-0 | POL | 41.72 |
| 7217 | Lm-2 | FRA | 53.38 |
| 7218 | Le-0 | NED | 25.27 |
| 7223 | Li-2:1 | GER | 53.05 |
| 7236 | Litva | LTU | 58.04 |
| 7244 | Mnz-0 | GER | 26.54 |
| 7250 | Me-0 | GER | 22.54 |
| 7255 | Mh-0 | GER | 16.05 |
| 7258 | Nw-0 | GER | 19.87 |
| 7268 | Np-0 | GER | 37.53 |
| 7276 | Ob-0 | GER | 4.02 |
| 7280 | Old-1 | GER | 29.41 |
| 7282 | Or-0 | GER | 15.20 |
| 7287 | Ove-0 | GER | 18.86 |
| 7298 | Pi-0 | AUT | 21.78 |
| 7305 | Pt-0 | GER | 34.12 |
| 7306 | Pog-0 | CAN | 37.61 |
| 7314 | Ragl-1 | UK | 10.33 |
| 7316 | Rhen-1 | NED | 62.80 |
| 7319 | Rome-1 | ITA | 47.29 |
| 7320 | Rou-0 | FRA | 28.26 |
| 7332 | Seattle-0 | USA | 47.93 |
| 7333 | Sei-0 | ITA | 42.03 |
| 7337 | Si-0 | GER | 28.05 |
| 7342 | Su-0 | UK | 41.12 |
| 7343 | Sp-0 | GER | 39.21 |
| 7344 | Sg-1 | GER | 53.69 |
| 7347 | Stw-0 | RUS | 40.64 |
| 7349 | Ta-0 | CZE | 70.45 |
| 7350 | Tac-0 | USA | 56.18 |
| 7353 | Tha-1 | NED | 36.96 |
| 7356 | Tol-0 | USA | 42.10 |
| 7372 | Tscha-1 | AUT | 38.95 |
| 7377 | Tul-0 | USA | 57.47 |
| 7378 | Uk-1 | GER | 37.05 |
| 7382 | Utrecht | NED | 33.57 |
| 7383 | Van-0 | CAN | 44.11 |
| 7384 | Ven-1 | NED | 37.85 |
| 7387 | Vind-1 | UK | 49.26 |
| 7394 | Wa-1 | POL | 25.28 |
| 7404 | Wc-1 | GER | 24.48 |
| 7411 | Wl-0 | GER | 27.05 |
| 7419 | Db-1 | GER | 69.10 |
| 7424 | Jl-3 | CZE | 74.25 |
| 7430 | Nc-1 | FRA | 40.40 |
| 7460 | Da(1)-12 | CZE | 40.01 |
| 7471 | RLD-1 | UNK | 34.48 |
| 7514 | RRS-7 | USA | 56.34 |
| 7520 | Lp2-2 | CZE | 55.43 |
| 7523 | Pna-17 | USA | 55.11 |
| 8214 | Gy-0 | FRA | 55.69 |
| 8264 | Bla-1 | ESP | 29.46 |
| 8311 | In-0 | AUT | 51.15 |
| 8312 | Is-0 | GER | 37.85 |
| 8337 | Mir-0 | ITA | 53.40 |
| 8343 | Na-1 | FRA | 43.97 |
| 8354 | Per-1 | RUS | 41.14 |
| 8357 | Pla-0 | ESP | 47.29 |
| 8366 | Rd-0 | GER | 12.86 |
| 8420 | Kelsterbach-4 | GER | 29.51 |
| 8424 | Kas-2 | IND | 48.16 |
| 9758 | Altai-5 | CHN | 70.36 |
| 9759 | Anz-0 | IRN | 40.11 |
| 9764 | Qar-8a | LBN | 60.15 |
| 9766 | Westkar-4 | KGZ | 59.77 |

**Supplementary Table 2.** List of SNPs significanlty associated with stained area (%) at 72 h after TZ staining and the nearest annotated gene. SNPs information was obtained from the GWA-Portal (https://gwas.gmi.oeaw.ac.at/). Score: -Log10 p-value, Maf: Minor allele frequency.

| Chr | Position | Score | Maf | Effect | Function | Codon | Amino acid substitution | Closest gene |
|---|---|---|---|---|---|---|---|---|
| 2 | 16969091 | 7.64 | 0.23 | Intergenic | - | - | - | AT2G40660 |
| 2 | 16969110 | 7.64 | 0.23 | Intergenic | - | - | - | AT2G40660 |
| 2 | 16969150 | 7.64 | 0.23 | Intergenic | - | - | - | AT2G40660/AT2G40670 |
| 2 | 16969193 | 7.64 | 0.23 | Intergenic | - | - | - | AT2G40660/AT2G40670 |
| 2 | 16969231 | 7.64 | 0.23 | Intergenic | - | - | - | AT2G40660/AT2G40670 |
| 2 | 16969241 | 7.64 | 0.23 | Intergenic | - | - | - | AT2G40660/AT2G40670 |
| 2 | 16969242 | 7.64 | 0.23 | Intergenic | - | - | - | AT2G40660/AT2G40670 |
| 2 | 16969245 | 7.64 | 0.23 | Intergenic | - | - | - | AT2G40660/AT2G40670 |
| 2 | 16969267 | 7.64 | 0.23 | Intergenic | - | - | - | AT2G40660/AT2G40670 |
| 2 | 16969277 | 7.64 | 0.23 | Intergenic | - | - | - | AT2G40660/AT2G40670 |
| 2 | 16969318 | 7.64 | 0.23 | Intergenic | - | - | - | AT2G40660/AT2G40670 |
| 2 | 16969414 | 8.89 | 0.24 | Intergenic | - | - | - | AT2G40660/AT2G40670 |
| 2 | 18431495 | 7.61 | 0.24 | Non synonymous coding | Missense | Tgt/Agt | C161S | AT2G44700 |
| 4 | 429928 | 7.65 | 0.40 | Synonymous coding | Silent | caT/caC | H546 | AT4G00990 |
| 5 | 19505864 | 9.07 | 0.35 | Intergenic | - | - | - | AT5G48110/AT5G48120 |

**Supplementary Table 3.** Composition of suberin aliphatics, phenolics and others in WT and *arr16* mutant seeds (µg/mg dry residue). Values presented are means ±SE (* $P < 0.05$, ** $P < 0.01$, *** $P < 0.001$, Dunnett's test)

| Monomer | WT | *arr16#1* | *arr16#2* |
|---|---|---|---|
| 18OH | 0.085±0.005 | 0.079±0.005 | 0.096±0.009 |
| 20:0-OH | 0.088±0.005 | 0.086±0.005 | 0.085±0.006 |
| iso22:0-OH | 0.033±0.002 | 0.028±0.001 | 0.033±0.001 |
| 22:0-OH | 0.361±0.020 | 0.336±0.015 | 0.329±0.020 |
| 20:0-diol | 0.056±0.005 | 0.066±0.002 | 0.070±0.005 |
| 22:0-diol | 0.284±0.014 | 0.278±0.008 | 0.302±0.019 |
| 16:0 | 0.136±0.004 | 0.150±0.014 | 0.160±0.009 |
| 18:0 | 0.035±0.002 | 0.040±0.010 | 0.038±0.003 |
| 18:1 | 0.056±0.003 | 0.084±0.012 | 0.081±0.004 ** |
| 18:2 | 0.114±0.015 | 0.158±0.007 * | 0.178±0.010 ** |
| 20:0 | 0.043±0.003 | 0.043±0.004 | 0.048±0.004 |
| iso22:0 | 0.017±0.002 | 0.020±0.007 | 0.014±0.001 |
| 22:0 | 0.087±0.003 | 0.077±0.005 | 0.092±0.008 |
| 24:0 | 0.277±0.014 | 0.260±0.011 | 0.286±0.015 |
| 26:0 | 0.123±0.005 | 0.148±0.007 * | 0.144±0.010 |
| 26:1 | 0.161±0.006 | 0.178±0.010 | 0.170±0.009 |
| 16:0-DCA | 0.122±0.004 | 0.146±0.007 * | 0.161±0.015 |
| 18:1-DCA | 0.393±0.009 | 0.434±0.015 * | 0.480±0.030 * |
| 18:2-DCA | 0.953±0.046 | 1.017±0.033 | 1.173±0.040 ** |
| 22:0-DCA | 0.241±0.008 | 0.270±0.015 | 0.291±0.028 |
| 23:0-DCA | 0.026±0.003 | 0.034±0.002 | 0.044±0.007 * |
| 24:0-DCA | 1.284±0.033 | 1.453±0.048 * | 1.648±0.129 * |
| 16:0-ωOH | 0.083±0.003 | 0.096±0.002 ** | 0.108±0.011 |
| 18:0-ωOH | 0.014±0.002 | 0.018±0.003 | 0.017±0.001 |
| 18:1-ωOH | 0.228±0.005 | 0.258±0.016 | 0.281±0.020 |
| 18:2-ωOH | 0.258±0.005 | 0.280±0.008 * | 0.324±0.011 ** |
| 18:0 diOH DCA | 0.282±0.010 | 0.349±0.060 | 0.388±0.030 |
| 18:1-triOH | 0.199±0.009 | 0.254±0.030 | 0.224±0.029 |
| 20:0-ωOH | 0.018±0.002 | 0.023±0.003 | 0.028±0.006 |
| iso22:0-ωOH | 0.105±0.010 | 0.092±0.004 | 0.099±0.006 |
| 22:0-ωOH | 0.461±0.005 | 0.451±0.011 | 0.481±0.040 |
| 23:0-ωOH | 0.042±0.002 | 0.052±0.005 | 0.050±0.006 |
| iso24:0-ωOH | 0.043±0.003 | 0.048±0.002 | 0.050±0.003 |
| 24:0-ωOH | 1.744±0.029 | 1.732±0.043 | 1.907±0.141 |
| 2OH-24:0 | 0.091±0.002 | 0.098±0.003 | 0.111±0.007 * |
| Unknown 1 | 0.416±0.009 | 0.382±0.059 | 0.463±0.029 |
| mz297 | 0.158±0.008 | 0.296±0.047 * | 0.273±0.023 ** |
| mz75/281 | 0.164±0.011 | 0.229±0.010 ** | 0.249±0.017 ** |
| Methyl dihydroxy benzoate | 0.606±0.048 | 1.095±0.044 *** | 1.284±0.141 ** |
| Ferulic acid | 0.492±0.013 | 0.533±0.029 | 0.576±0.050 |
| Caffeic acid | 0.097±0.006 | 0.093±0.005 | 0.104±0.008 |

**Supplementary Table 4.** List of primers used for genotyping of T-DNA insertion lines.

| Line | Gene/T-DNA border | Forward primer | Reverse primer |
|---|---|---|---|
| *salk_020583* | *AT5G13930 (TT4)* | TCGAATAGACCTGTCCAGCAC | CTTCTCTGGACACCAGACAGG |
| | T-DNA border | ATTTTGCCGATTTCGGAAC | - |
| *salk_142105* | *AT2G40670 (ARR16)* | CAAGATCCTGAAACAGATCCG | TTCCAGGCATACAGTAATCGG |
| | T-DNA border | ATTTTGCCGATTTCGGAAC | - |
| *salkseq_112294.2* | *AT2G40670 (ARR16)* | ACCAATCTTTTGCGGATGAC | TAACCATTGAAAATGGAGGCC |
| | T-DNA border | ATTTTGCCGATTTCGGAAC | - |
| *ahk2-5* | *AT5G35750 (AHK2)* | GCAAGAGGCTTTAGCTCCAA | TTGCCCGTAAGATGTTTTCA |
| | T-DNA border | - | GCCTTTTCAGAAATGGATAAATAGCCTTGCTTCC |
| *ahk3-7* | *AT1G27320 (AHK3)* | CCTTGTTGCCTCTCGAACTC | CGCAAGCTATGGAGAAGAGG |
| | T-DNA border | CCCATTTGGACGTGTAGACAC | - |
| *cre1-2* | *AT2G01830 (AHK4)* | CTCTTTTGTTCTTGAATTCGC | ATCCTGCAACATTCTAGCTC |
| | T-DNA border | - | ATAACGCTGCGGACATCTAC |

**Supplementary Table 5.** List of primers used for RT-qPCR.

| Gene name | AGI gene code | Forward primer | Reverse primer |
|---|---|---|---|
| *KCS2* | AT1G04220 | GGTTGAACCTGAAGAAGC | CCGACGCCTTCGTTATCG |
| *ABCG2* | AT2G37360 | GGAGACTTGTGTGACGACGG | TCACTTCCGTTTGTTCTTGCTACC |
| *GPAT5* | AT3G11430 | TCCGGATCTTTCTTGGAGCCGT | GGTTCTGTGAGTACACACAAAGAGCA |
| *FAR4* | AT3G44540 | ATTCATATTCCCGGCCTCA | TCGGTCCAAGCAAATTTTATG |
| *FAR5* | AT3G44550 | GGTTTCCTTGCGAAGGTTTT | GTGGCAGCTTCATTGTCAGA |
| *ABCG20* | AT3G53510 | TCGGGCTATGGATATTGCGG | ACACGAGAAGACGGAACCG |
| *ABCG6* | AT5G13580 | GACTCCACGCGTCGTAGTC | TCCGGCGAACTTTGACGC |
| *FAR1* | AT5G22500 | CCTCATCACCCATGTGCTTA | CACGCCAACACAAACTGTAAA |
| *CYP86B1* | AT5G23190 | AGTGACCCTCTGGTTTCACG | CTTCTGACGCAAGGCATGTA |
| *ASFT* | AT5G41040 | TCAAAAGGAACCAGCTTTGG | TTCCCTCTCTCCTCGGATTT |
| *CYP86A1* | AT5G58860 | CTCCGTTACCGGGTTTTTC | TCACCACGAGGTTGCAAATA |
| *FACT* | AT5G63560 | GGTTGTTCAGGTGACAAATTTCAA | CGATACCATCGAACATATTGTGGTT |
| *EF1α* | AT5G60390 | CTTCTTGCTTTCACCCTTGGTGT | TGTCAGGGTTGTATCCGACCTT |
| N.A. | AT4G12590 | GAGATGAAAATGCCATTGATGAC | GCACCCAGACTCTTTGATG |
